# Taxon-specific or universal? Using target capture to study the evolutionary history of rapid radiations

**DOI:** 10.1101/2021.05.20.444989

**Authors:** Gil Yardeni, Juan Viruel, Margot Paris, Jaqueline Hess, Clara Groot Crego, Marylaure de La Harpe, Norma Rivera, Michael H. J. Barfuss, Walter Till, Valeria Guzmán-Jacob, Thorsten Krömer, Christian Lexer, Ovidiu Paun, Thibault Leroy

## Abstract

Target capture has emerged as an important tool for phylogenetics and population genetics in non-model taxa. Whereas developing taxon-specific capture probes requires sustained efforts, available universal kits may have a lower power to reconstruct relationships at shallow phylogenetic scales and within rapidly radiating clades. We present here a newly-developed target capture set for Bromeliaceae, a large and ecologically-diverse plant family with highly variable diversification rates. The set targets 1,776 coding regions, including genes putatively involved in key innovations, with the aim to empower testing of a wide range of evolutionary hypotheses. We compare the relative power of this taxon-specific set, Bromeliad1776, to the universal Angiosperms353 kit. The taxon-specific set results in higher enrichment success across the entire family, however, the overall performance of both kits to reconstruct phylogenetic trees is relatively comparable, highlighting the vast potential of universal kits for resolving evolutionary relationships. For more detailed phylogenetic or population genetic analyses, e.g. the exploration of gene tree concordance, nucleotide diversity or population structure, the taxon-specific capture set presents clear benefits. We discuss the potential lessons that this comparative study provides for future phylogenetic and population genetic investigations, in particular for the study of evolutionary radiations.

## 1 Introduction

Targeted sequencing approaches have emerged as a promising tool for studying evolutionary relationships in non-model taxa, enabling researchers to retrieve large data sets while requiring few genomic resources (Bossert & Danforth, 2018; Escudero, Nieto-Feliner, Pokorny, Spalink, & Viruel, 2020; McDonnell et al., 2021; Soto-Gomez et al., 2019). Using custom baits, the method largely retrieves the same loci across a wide taxonomic scale, obtains comparable and mergeable data sets, and may be combined with genome-skimming (E. M. Lemmon & Lemmon, 2013; Weitemier et al., 2014). Pre-existing knowledge of the targeted loci further provides opportunities to address specific questions on both deep and shallow timescales (Hale, Gardner, Viruel, Pokorny, & Johnson, 2020; A. R. Lemmon, Emme, & Lemmon, 2012). Finally, the method does not necessarily require a reference genome, is highly cost-effective, and with the ability to sequence herbarium samples, reduces the need for extensive sampling campaigns (Blaimer, Lloyd, Guillory, & Brady 2016; Hale et al. 2020; Weitemier et al., 2014). Target capture has been successfully applied to resolve phylogenies in diverse groups, from arthropods such as bees (*Xylocopa*, Blaimer et al., 2016; Apidae, Bossert et al., 2019) and Araneae (Hexathelidae, Hedin, Derkarabetian, Ramírez, Vink, & Bond, 2018) to mammals (Cetacea, McGowen et al., 2020), and in numerous plant groups (*Heuchera*, Folk, Mandel, & Freudenstein, 2015; Gesneriaceae, Ogutcen et al., 2021; Zingiberales, Sass, Iles, Barrett, Smith, & Specht, 2016 to name a few). The method’s utility for studies at micro-evolutionary scales has been to date marginally explored, but several studies have pointed to the ability to analyze genomic diversity and estimate population genomic parameters (Choquet et al., 2019; Christmas, Biffin, Breed, & Lowe, 2017; de La Harpe et al., 2019; Derrien & Ramos-Onsins, 2020; Sanderson, DiFazio, Cronk, Ma, & Olson, 2020). Nonetheless, the development of probes for target enrichment may pose several challenges: first, the need to identify regions conserved enough to ensure recovery, yet polymorphic enough to provide ample information (Soto-Gomez et al., 2019; Villaverde et al., 2018). Second, probe design requires detecting regions without pervasive copy number polymorphism (Kadlec, Bellstedt, Maitre, & Pirie, 2017; A. R. Lemmon et al., 2012), a particular challenge for angiosperms and other groups, where duplication events are ubiquitous (Van de Peer, Mizrachi, & Marchal, 2017).

In contrast, universal kits offer an attractive alternative that require reduced efforts to establish, and provide comparable data sets across wider ranges of taxa (Johnson et al., 2019; Kadlec et al., 2017). Such kits were designed to retrieve single-copy markers, for example, in the broad scope of amphibians (Hime et al., 2021), anthozoans (Quattrini et al., 2018), vertebrates (A. R. Lemmon et al., 2012) or angiosperms (Johnson et al., 2019). In the latter example, the Angiosperms353 kit is designed to target 353 single-copy genes across angiosperms. So far the kit has been employed successfully in resolving phylogenies, including but not limited to *Nepenthes* (Murphy et al., 2020), *Schefflera* (Shee, Frodin, Cámara-Leret, & Pokorny, 2020) and the rapid radiations of *Burmeistera* (Bagley, Uribe-Convers, Carlsen, & Muchhala, 2020) and *Veronica* (Thomas et al., 2021), establishing the kit as an eminent tool in macroevolutionary research. Its utility at microevolutionary levels is yet to be fully realized, although several works have established its suitability to deliver informative signals at a lower taxonomic level (Beck et al., 2021) and in acquiring population genomics parameters (Slimp, Williams, Hale, & Johnson, 2021). The use of highly-conserved markers in a universal kit may, however, limit resolution power. Generally, taxon-specific baits are expected to deliver a higher information content and hence more accurate results (Kadlec et al., 2017), as enrichment success is known to drop with the level of divergence between sequences used for probe design and the targeted taxa (Liu et al., 2019). However, one study comparing the power of the universal Angiosperms353 kit and a taxon-specific kit to resolve phylogenomic relationship in Cyperaceae reported surprisingly similar performance (Larridon et al., 2020) and similar findings were reported in Malinae (Ufimov et al., 2021) and in Ochnaceae (Shah et al., 2021). It remains to be established whether these findings apply to other taxa and other evolutionary scales, including at population level, where ample genomic variability is required to resolve intra-specific relationships and investigate patterns of genetic differentiation.

Until recently, the technology available to investigate evolutionary questions in rapidly evolving groups featuring high net diversification rates has presented major obstacles, in particular for non-model groups. Decreasing costs of sequencing coupled with an ever-growing plethora of bioinformatic tools for data processing and downstream analysis has led to an increase in the use of methods like whole-genome sequencing, RNA sequencing and restriction site associated DNA sequencing (RAD-Seq) in lieu of traditional methods employing few conserved markers (de La Harpe et al., 2017; McKain, Johnson, Uribe-Convers, Eaton, & Yang, 2018; Weitemier et al., 2014; Zimmer & Wen, 2013). Whole-genome sequencing however remains costly, posing barriers for research targeting large numbers of samples, organisms with large genomes and non-model organisms, for which the availability of high-quality genomic resources is often limited (Hollingsworth, Li, van der Bank, & Twyford, 2016; Supple & Shapiro, 2018). While RAD-seq is an affordable alternative and widely used in population genetics, the resulting data sets may fall short when screened for homologous sequences across distantly related lineages (but see e.g., Heckenhauer, Samuel, Ashton, Abu Salim, & Paun, 2018). Additionally, RAD-seq is less feasible when using degraded DNA from herbarium samples, and the use of short and inconsistently-represented loci across phylogenetic sampling may result in low information content and difficulties in assessing paralogy (E. M. Lemmon & Lemmon, 2013; McKain et al., 2018; Jones & Good, 2016).

Rapid evolutionary radiations are key stages in the evolutionary history across the Tree of Life and highly recurrent, hence an essential part of biodiversity research (Gavrilets & Losos, 2009; Givnish et al., 2014; Hughes, Nyffeler, & Linder, 2015; Soltis, Folk, & Soltis, 2019; Soltis & Soltis, 2004). Fast evolving groups provide potent opportunities to investigate important questions in evolutionary biology, such as the interplay between ecological and evolutionary processes in shaping biodiversity. A few notable study systems are the cichlid fish (McGee et al., 2020; Salzburger, 2018), *Heliconius* butterflies (Dasmahapatra et al., 2012; Moest et al., 2020), *Anolis* lizards (McGlothlin et al., 2018; Stroud & Losos, 2020), Darwin’s finches (Lamichhaney et al., 2015; Zink & Vázquez-Miranda, 2019), white-eyes birds (Moyle, Filardi, Smith, & Diamond, 2009) and New World lupins (Nevado, Atchison, Hughes, & Filatov, 2016). Nevertheless, much remains unknown about the genomic basis underlying species diversification outside these intensively studied systems.

Research of rapidly diversifying lineages presents several challenges. First, a brief diversification period typically leads to imperfect reproductive barriers and incomplete lineage sorting, reflected in significant gene tree discordance and ambiguous relationships (Degnan & Rosenberg, 2009; Lamichhaney et al., 2015; Pease, Haak, Hahn, & Moyle, 2016; Straub et al., 2014). In addition, understanding ‘speciation through time’ poses a methodological challenge, and requires connecting two conceptual worlds: macroevolutionary investigations, concerned with spatial and ecological patterns over deeper timescales, and microevolutionary approaches, providing insight into the processes acting during population divergence and speciation (Bragg, Potter, Bi, & Moritz, 2016; de La Harpe et al., 2017). Resolving phylogenomic relationships and disentangling the contribution of different genomic processes through time typically requires large-scale genomic datasets and thorough taxon sampling efforts (E. M. Lemmon & Lemmon, 2013; Linder, 2008; Straub et al., 2012).

Here, we present Bromeliad1776, a new bait set for targeted sequencing, designed to address a wide range of evolutionary hypotheses in Bromeliaceae: from producing robust phylogenies to studying the interplay of genomic processes during speciation and the genetic basis of trait shifts, such as photosynthetic and pollination syndrome. This highly diverse Neotropical radiation provides an excellent research system for studying the drivers and constraints of rapid adaptive radiation (Benzing, 2000; Givnish et al., 2011; Loiseau et al., 2021; Mota et al., 2020; Palma-Silva & Fay, 2020; Wöhrmann, Michalak, Zizka, & Weising, 2020). Bromeliaceae as a whole is considered an adaptive radiation (Benzing, 2000; Givnish et al., 2011) and contains several rapidly radiating lineages, most notably within Bromelioideae (Aguirre-Santoro, Salinas, & Michelangeli, 2020) and Tillandsioideae (Loiseau et al., 2021). It is a species-rich and charismatic monocot family, consisting of over 3,000 species, including crops in the genus *Ananas* and other economically important species (Luther, 2008). Members of the family are characterized by a distinctive leaf rosette that often impounds rainwater in central tanks (phytotelmata). A diversity of arthropods and other animal species and microbes reside in bromeliad tanks, in some cases even leading to protocarnivory and other forms of nutrient acquisition (Givnish, Burkhardt, Happel, & Weintraub, 1984; C. Leroy, Carrias, Céréghino, & Corbara, 2016). Bromeliads present a diversity of repeatedly evolving adaptive traits, which allowed them to occupy versatile habitats and ecological niches (Benzing, 2000). CAM photosynthesis, water-absorbing trichomes, formation of tank habit, extensive rates of epiphytism and a diversity of pollination syndromes are some of the adaptations correlated with high rates of diversification within the family (Benzing, 2000; Crayn, Winter, & Smith, 2004; Givnish et al., 2014; Kessler, Abrahamczyk, & Krömer, 2020; Quezada & Gianoli, 2011).

To assess the utility of the Bromeliad1776 kit, we performed a comparison between our taxon-specific kit and the universal Angiosperms353 kit using several methods across different evolutionary time-scales. We present Bromeliad1776 in the light of methodological considerations on bait design, data handling, analyses and other practical considerations.

## 2 Materials and Methods

### 2.1 Custom bait design

Whole-genome sequences and gene models from *Ananas comosus* v.3 (Ming et al., 2015) were used to design a bait set aiming to target i) single-copy protein coding genes distributed across the whole genome, ii) genes previously described as associated with key innovation traits in Bromeliaceae (see below), iii) markers previously used for phylogenomic inference in Bromeliaceae and iv) genes orthologous to those in the Angiosperms353 bait set. The 1776 selected genes are detailed in Supporting information Table S1.

Genes in subset *i* were selected based on genetic diversity parameters calculated using whole-genome sequence and RNAseq data previously published by de La Harpe et al., (2020; data publicly available online at SRA Bioproject PRJNA649109) with the PopGenome R package v.2.1.6 (Pfeifer, Wittelsbürger, Ramos-Onsins, & Lercher, 2014). Genomic regions were retained in this category if they shared at least 70% identity between *A. comosus* and *T. sphaerocephala*, and if they had nucleotide diversity (*π*) values not exceeding the 90% quantile of the (*π*) distribution across genes for four *Tillandsia* species (*Tillandsia australis*, *Tillandsia fasciculata*, *Tillandsia floribunda* and *T. sphaerocephala*; data and analysis performed by de La Harpe et al. (2020). We further excluded genes with a total exonic size smaller than 1,100 bp, or individual exons smaller than 120 bp. Next, copy-number variation was calculated based on clustering of *A. comosus* and *Tillandsia* transcriptome assemblies to generate three copy number categories - “single copy”, “low copy” (i.e., less than five copies) and “high copy” (i.e., five or more copies). We included only single-copy genes in the design for bait subset *i*. Finally, we excluded genes that were located in genomic regions outside those assigned to linkage groups in the *A. comosus* reference (Ming et al., 2015). A total of 1,243 genes were identified for this part.

The bait subset of genes associated with key innovative traits in Bromeliaceae (subset *ii* above) included (1) genes putatively under positive selection along branches relevant to C3/CAM shifts (de La Harpe et al., 2020), (2) genes that exhibit differential gene expression between CAM and C3 *Tillandsia* species (de La Harpe et al., 2020) and (3) genes putatively associated with photosynthetic and developmental functions, or with flavonoid and anthocyanin biosynthesis, according to the literature (e.g. Ming et al., 2015; Palma-Silva, Ferro, Bacci, & Turchetto-Zolet, 2016; Wai et al., 2017; Goolsby, Moore, Hancock, Vos, & Edwards, 2018). *Ananas comosus* genes with the highest match scores (calculated as lowest E-score in BLASTP, Madden (2013) against the sequences of genes from the literature were added to the bait set (see Supporting information Table S2 for details). A total of 1,612 genes underpinning innovative traits were included in the bait design, regardless of criteria used for subset *i* for size, similarity and duplication rate.

Markers previously used for phylogenomic inference in Bromeliaceae (subset *iii*) were obtained from the literature, spanning 13 genes (e.g. Barfuss et al., 2016; Machado et al., 2020; Schulte, Barfuss, & Zizka, 2009, see TS2 for full list). Genes orthologous to those in the Angiosperms353 bait set (Johnson et al., 2019) were identified using the orthologous gene models from *A. comosus* based on gene annotations (Ming et al., 2015) or using BLASTP (Madden, 2013), totalling 281 genes.

Finally, we used a draft genome of *T. fasciculata* (Jaqueline Hess, personal communication) to exclude from all candidates genes that exhibited multiple BLASTN hits, if they have not been previously described as duplicated within the genus (de La Harpe et al., 2020). Specifically, we excluded genes that matched another genomic sequence of at least 100bp with high similarity score (*>* 80%) and low E-value (*<* 10*^−^*^5^). In an additional round of filtering performed by the manufacturer of the final bait set, Arbor Biosciences (Ann Arbor, MI, USA), multi-copy genes with sequences that are more than 95% identical were collapsed into a single sequence and baits with more than 70% GC content or containing at least 25% repeated sequences were excluded. In addition, targets including exons smaller than 80 bp were completed with regions flanking the exons according to the *A. comosus* reference genome. The final kit included 1776 genes: 801 genes in subset *i*, 681 genes associated with key innovative traits, 13 genes representing phylogenetic markers and 281 genes orthologous to the Angiosperms353 set. Probes were designed with 57,445 80-mer baits tiling across targets in 2x coverage, targeting approximately 2.3Mbp. The kit is subsequently referred to as the Bromeliad1776 bait set. Further specifications can be found in Supporting information Tables S1 and S2 and in the github repository: https://github.com/giyany/Bromeliad1776/tree/main/MS_2021_scripts.

### 2.2 Plant material collection

We sampled a total of 70/72 Bromeliaceae samples (for Angiosperms353 and for Bromeliad1776, accordingly; Supporting information Table S3), including 56 accessions from the Tillandsioideae subfamily and 16 representing the other subfamilies, except Navioideae. The divergence time between Tillandsioideae and subfamily Bromelioideae to which *A. comosus* belongs is estimated at 15Mya (according to Givnish et al. 2014). Within Tillandsioideae, we sampled 38/40 individuals from five species of the *Tillandsia* subgenus *Tillandsia* (‘clade K’ in Barfuss et al. (2016); Sampling in Mexican populations illustrated in Supporting information Figure S1).

### 2.3 Library preparation & enrichment

DNA extractions were performed using a modified CTAB protocol (Doyle & Doyle, 1987), purified using Nucleospin^®^ gDNA cleanup kit from Macherey-Nagel (Hudlow et al., 2011) following the supplier’s instructions with a two-fold elution step and finally quantified with Qubit^®^ 3.0 Fluorometer (Life Technologies, Ledeberg, Belgium).

For each sample, 200ng DNA was sheared using Bioruptor^®^ Pico sonication device (Diagenode, Seraing, Belgium) aiming for an average insert size of 350bp, dried in a speed vacuum Eppendorf concentrator 5301 (Eppendorf, Germany) and eluted in 30 L ddH_2_O. Genomic libraries were prepared using the NEBNext^®^ Ultra TM II DNA Library Prep Kit for Illumina® (New England Biolabs, Ipswich, MA, United States) using reagents at half volumes following Hale et al. (2020) and using 11 PCR cycles, increased up to 13 cycled for libraries with low genomic output. Samples were double-indexed with NEBNext^®^ Multiplex Oligos for Illumina^®^ (New England Biolabs, Ipswich, MA, USA). Fragment sizes were inspected with Agilent Bioanalyzer (Agilent Technologies, Santa Clara, CA, USA) and concentrations were measured with Qubit^®^ 3.0 Fluorometer. Subpools of 11-14 equimolar genomic libraries were prepared using phylogenetic proximity and DNA concentrations of the genomic libraries, which ranged from 2.62 to 118.0 ng/ L, following Soto-Gomez et al. (2019).

We used the Angiosperms353 and the Bromeliad1776 bait sets from Arbor Biosciences (Ann Arbor, MI, USA) to enrich each subpool of genomic libraries independently with a single hybridization reaction of myBaits^®^ target capture kits from Arbor Biosciences (Ann Arbor, MI, USA), following Hale et al. (2020). Average fragment size and DNA yield were estimated for each subpool using Agilent Bioanalyzer and Qubit^®^ 3.0 Fluorometer. Subpools were then pooled in equimolar conditions and sequenced at Vienna BioCenter Core Facilities (Vienna, Austria) on Illumina^®^ NextSeq^TM^ 550 (2×150bp, Illumina, San Diego, CA). Sequencing was conducted independently for either bait kit.

### 2.4 Data processing

The raw sequence data in BAM format was demultiplexed using deML v.1.1.3 (Renaud, Stenzel, Maricic, Wiebe, & Kelso, 2015) and samtools view v.1.7 (Li et al., 2009), converted to fastq using bamtools v.2.4.0 (Barnett, Garrison, Quinlan, Strömberg, & Marth, 2011) and quality checked using FastQC v.0.11.7 (Andrews, 2010). Reads were then trimmed for adapter content and quality using TrimGalore v.0.6.5 (Krueger, 2019), a wrapper tool around FastQC and Cutadapt, using settings –fastqc –retain unpaired. Sequence quality and adapter removal was confirmed with FastQC reports.

Quality and adapter-trimmed reads were aligned to *A. comosus* reference genome v.3 (Ming et al., 2015) using bowtie2 (Langmead & Salzberg, 2012) with the –very-sensitive-local option to increase sensitivity and accuracy. Samtools (Li et al., 2009) was then used to remove low quality mapping and sort alignments by position, and PCR duplicates were marked using MarkDuplicates from PicardTools v.2.25 (*Picard Toolkit*, 2019). Summary statistics of the mapping step were generated using samtools stats. Variants were called using freebayes v1.3.2-dirty (Garrison & Marth, 2012) and sites marked as MNP/complex were decomposed and normalized using the script ‘vcfallelicprimitives’ from vcflib (Garrison, 2012). Next, AN/AC field was calculated using bcftools v.1.7 (Li, 2011) and variant calls were filtered using vcflib (Garrison & Marth, 2012) and bcftools. Given that freebayes does not perform automatic variant filtering steps, we identified sets of parameters that generate reliable final SNP sets, based on two independent criteria: the highest transition/transversion ratios as reported by SnpSift (SnpEff suite, Cingolani et al., 2012) and the lowest *π*_N_/*π*_S_ (see section 2.7 below). After a detailed evaluation, we used the following criteria to generate two high quality SNP sets, one for each bait-set: we considered genotype calls with per-sample coverage below 10× as missing (NA) and excluded variants (i) marked as indels or neighboring indels within a distance of 3 bp, (ii) with depth of coverage at the SNP level lower than 500×, (iii) with less than ten reads supporting the alternate allele at the SNP level, or (iv) with more than 40% missing data. All genes in the Bromeliad1776 that passed the filtering criteria were included in the SNP set, regardless of their function. Summary statistics of the final SNP sets were generated using the script vcf2genocountsmatrix.py, namely the total number of SNPs, the proportion of on-target SNPs and the proportion of SNPs in some specific genomic contexts, with *A. comosus* genome v.3 as a reference. The full data processing script align_and_trim.sh and the vcf2genocountsmatrix.py script are both available at https://github.com/giyany/Bromeliad1776.

### 2.5 Bait specificity and efficiency

To explore bait specificity, we calculated the percentage of high quality trimmed reads on-target using samtools stats and bedtools intersect v2.25.0 (Quinlan & Hall, 2010) using the script calculat_bait_target_specifity.sh (available from https://github.com/giyany/Bromeliad1776). Targets for Bromeliad1776 were defined as the bait sequences plus their 500 bp flanking regions. Targets for Angiosperms353 were defined using orthogroups to *A. comosus*: gene annotations from the bait set were used to assign genes to orthogroups using OrthoFinder (Emms & Kelly, 2019). When several orthogroups were found for a single Angiosperms353 gene, we included all, resulting in 559 *A. comosus* genes assigned to orthogroups. Within the orthgroups, targets were again defined as exonic regions plus their 500 bp flanking regions.

To provide insights into determinants of bait capture success, we calculated bait efficiency for all baits of Bromeliad1776. For each bait, efficiency was calculated as the number of high-quality reads uniquely mapping to each bait target region, averaged over samples. We then tested for the correlation of capture efficiency to several bait characteristics (copy number, GC content, number and size of exons in targeted gene, size of baits and phylogenetic distance to *A. comosus*) with a generalized linear model or Kruskal-Wallis test in R v.4.0.3 (R Core Team, 2020) using a negative binomial family.

### 2.6 Phylogenomic analyses

We inferred phylogenomic relationships for all samples using two methods: a concatenation method, and a coalescent-based species tree estimation. The latter method was included as concatenation methods do not account for gene tree incongruence, which may result in high support for an incorrect topology (Kubatko & Degnan, 2007), especially in the presence of notable incomplete lineage sorting. In addition, gene tree incongruence analysis provides insight into molecular genome evolution, including the extent of incomplete lineage sorting and other genomic processes such as hybridization and introgression (Galtier & Daubin, 2008; Wendel & Doyle, 1998).

We used the the variant and non-variant genotypes to create a phylip matrix with vcf2phylip v.2.0 (Ortiz, 2019) and constructed a maximum-likelihood species tree for each bait set with RAxML-NG v.0.9.0 (Kozlov, Darriba, Flouri, Morel, & Stamatakis, 2019), using 250 bootstrap replicates and a GTR model with an automatic MRE-based bootstrap convergence test. Next, we constructed a species tree using ASTRAL-III v.5.7.7 (hereafter: ASTRAL, Zhang, Rabiee, Sayyari, & Mirarab, 2018). For both the Angiosperms353 and the Bromeliad1776 sets, we separated the matrix into independent genomic windows, defining each window as a gene according to the known exons and a 500bp flanking region. For Angiosperms353, we extracted the 559 genes (assigned to orthogroups as explained above) as genomic windows using bedtools intersect. For Bromeliad1776, genomic windows were extracted using the *A. comosus* gene sequences included in bait design. All loci and all accessions were included in species tree inference regardless of the percentage of missing data, since taxon completeness of individual gene trees is important for statistical consistency of this approach, and we expected only low levels of fragmentary sequences (Mirarab, 2019; Nute, Chou, Molloy, & Warnow, 2018). After excluding genes with zero coverage, 269 genes and 1,600 genes were included in species tree inference for Angiosperms353 and Bromeliad1776, respectively.

For each gene, a maximum-likelihood gene tree was inferred using ParGenes (Morel, Kozlov, & Stamatakis, 2019) with RAxML-NG (Kozlov et al., 2019), using a GTR model with an automatic MRE-based bootstrap convergence test. Loci with insufficient signal may reduce the accuracy of species tree estimation (Mirarab, 2019), hence, in all gene trees, nodes with a bootstrap support smaller than ten were collapsed using Newick utilities (Junier & Zdobnov, 2010). A species tree was then generated in ASTRAL with quartet support and posterior probability for each tree topology. The number of conflicting gene trees was calculated using phyparts and visualized using the script phypartspiecharts.py (available from https://github.com/mossmatters/MJPythonNotebooks).

### 2.7 Population structure and nucleotide diversity estimates

To explore the genetic structure within the *Tillandsia* species complex, we focused on five species from 15 localities (Supporting information Table S3 and Supporting information Figure S1). We first used plink v.1.9 (Chang et al., 2015) to filter out SNPs in linkage disequilibrium. Population structure was further explored through individual ancestry analysis, with identity-by-descent matrix calculated by plink and inference of population structure using ADMIXTURE v.1.3. with K values ranging from one to ten, and 30 replicates for each K, using a block optimization method (Alexander & Lange, 2011). A summary of the ADMIXTURE results was obtained and presented using pong (Behr, Liu, Liu-Fang, Nakka, & Ramachandran, 2016). The set of LD-pruned biallelic SNPs was further filtered to allow a maximum of 10% missing data and used to perform a principal components analysis (PCA) with SNPRelate v.1.20.1 (Zheng et al., 2012). Finally, for each *Tillandsia* species, we used the strategy of T. Leroy et al. (2021) to compute synonymous (*π*_S_) and non-synonymous (*π*_N_) nucleotide diversities and Tajima’s D, from fasta sequences using seq_stat_coding (T. Leroy et al., 2021).

## 3 Results

### 3.1 Higher mapping rates and capture efficiency for taxon-specific set

On average, 4,401,958 (803,464-12,693,516) paired-end reads per accession were generated per Angiosperms353 library and 2,962,023 (1,282,762-6,298,880) per Bromeliad1776 library. Overall, the mapping rates to the *A. comosus* reference genome were higher for libraries enriched with Bromeliad1776, with an average mapping rate of 82.3% (61.8%-95.9%) and 42.8% (22.1%-77.9%), for Bromeliad1776 and Angiosperms353, respectively (Supporting information Figure S2, Supporting information Table S4). Higher mapping rates were recorded for subfamilies Bromelioideae and Puyoideae, as compared to Tillandsioideae, for both the Angiosperms353 and Bromeliad1776 sets (see Supporting information Figures S3 and S4, respectively). This may reflect the effect of reference bias, and in the case of Bromeliad1776, it may be further amplified by our kit design based on *A. comosus* (subfamily Bromelioideae). Bait specificity was high for Bromeliad1776 with on average 90.4% reads on-target (76.5%-94.2%), while for Angiosperms353 bait specificity was 14.0% (4.6%-30.1%; see Supporting information Figure S2). Mapping rates and bait specificity were positively correlated for both bait sets (GLM, P*<*0.01).

### 3.2 Bait efficiency depends on the genomic context

We investigated factors that may influence bait efficiency, starting with the contribution of gene copy number variation. We assumed three categories regarding the number of paralogs per orthogroup: single copy, low-copy (i.e., less than five copies) and high-copy (i.e., five or more copies). The number of gene copies had a significant effect on bait efficiency and post-hoc Dunn’s test supported significant differences in efficiency for comparisons between low-copy and high-copy, and between single-copy and low-copy (P=2.8*^−^*^44^). Low-copy genes exhibit the lowest enrichment success, suggesting that the bait efficiency is not simply correlated to the number of gene copies (Figure 1). We also recovered a significant effect of the intragenic GC content and GC content of the baits on bait efficiency (GLM, P=1.5*^−^*^68^). Finally, we investigated the possible link between efficiency and gene structure. Average exon sizes (P*<* 2.0*^−^*^16^) and total number of exons per gene (P=1.1*^−^*^89^) were also positively correlated with enrichment success. The size of the smallest exon for all targeted genes was however not correlated with bait efficiency. Sequence similarity, measured as percent of identity between Tillandsia sequences and those of *A. comosus*, was positively correlated with capture success (P=4.8*^−^*^13^; Figure 1).

**Figure 1.**
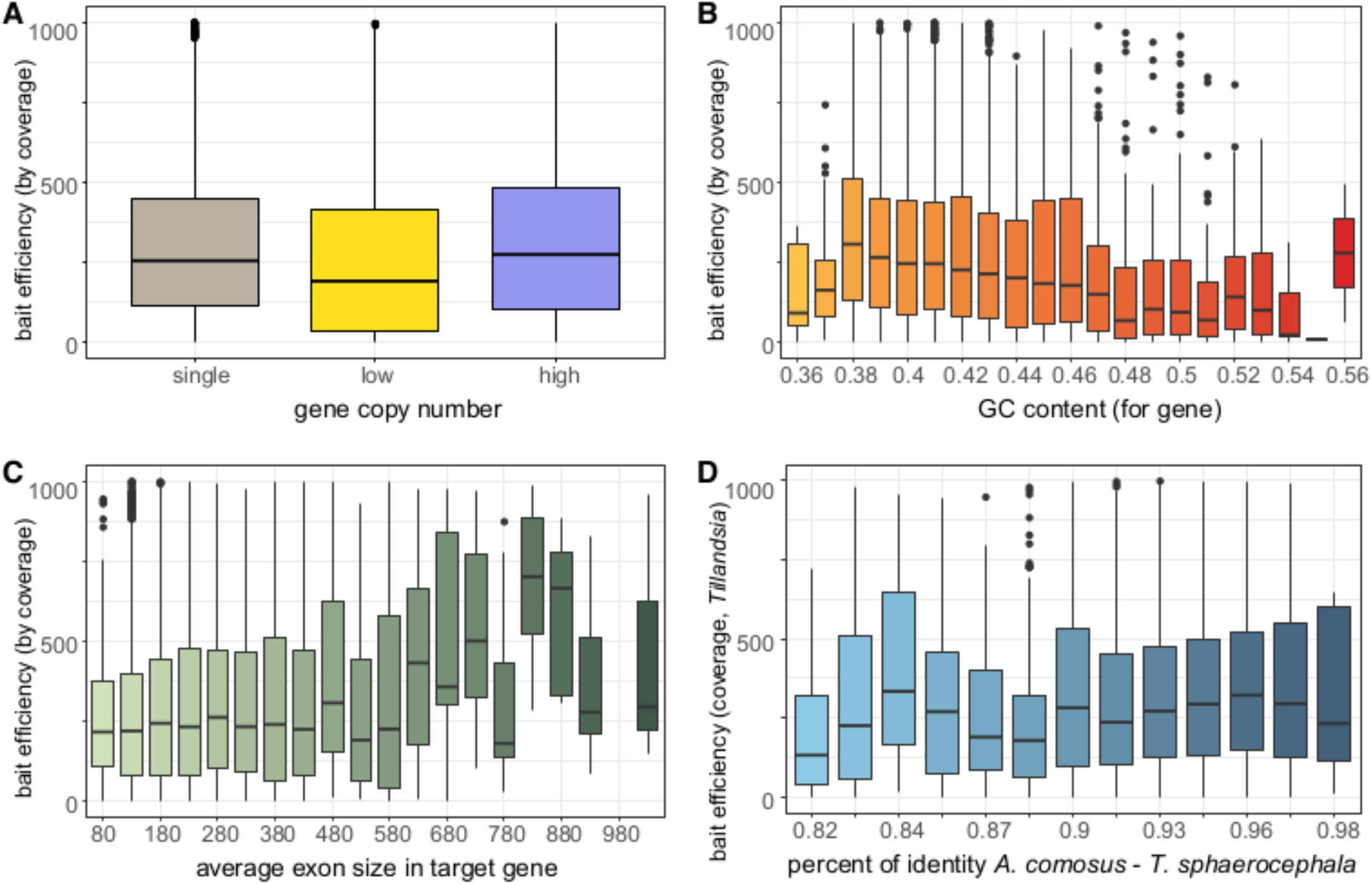
Effects of (A) putative gene copy number, (B) gene GC content, (C) average exon size, and (D) percent of identity on bait efficiency in Bromeliad1776 bait set, measured as the number of high-quality reads uniquely mapping to bait target region across samples. Continuous variable was binned and y-values higher than 1,000 excluded for visualization in B-D.

### 3.3 Both kits provided a large number of SNPs

After variant calling and filtering, we identified 47,390 and 209,186 high-quality SNPs for the Angiosperms353 and the Bromeliad1776 bait sets, respectively. On average, missing data represented 23.7% of genotype calls per individual in Angiosperms353, but only 6.3% for the Bromeliad1776 kit. The differences in amount of missing data are likely associated with the higher mean depth per site across the Bromeliad1776 kit (6,602), as compared to Angiosperms353 (3,437). Focusing on the subgenus *Tillandsia*, we identified 15,622 SNPs for Angiosperms353 (including a total of 18.9% missing data) compared to 65,473 polymorphic sites (2.9% missing data) for Bromeliad1776. In both full data sets and the subset including only *Tillandsia* samples, Bromeliad1776 recovered more variants in intronic regions compared with Angiosperms353. Angiosperms353 recovered a large proportion of off-target SNPs, whereas in Bromeliad1776 approximately 15% of the SNPs were recovered from flanking regions (Table 1). We discuss ascertainment bias that may rise due to the non-random selection of markers in the supporting information.

**Table 1.** Number and characteristics of the variants obtained for Angiosperms353 and Bromeliad1776.

|  | indv Nr. | SNP Nr. | site<br>mean depth | SNPs<br>in exonic<br>regions | SNPs<br>in intronic<br>regions | SNPs<br>in intergenic<br>regions | on-target<br>SNPs | flanking<br>SNPs | off-target<br>SNPs |
| --- | --- | --- | --- | --- | --- | --- | --- | --- | --- |
| <u>intragenic vcf</u> |  |  |  |  |  |  |  |  |  |
| Angiosperms353 | 70 | 47,390 | 3447 | 40,628<br>(85.7%) | 4,376<br>(9.2%) | 2,386<br>(5.1%) | 8,424<br>(17.8%) | 3,488<br>(7.4%) | 35,478<br>(74.8%) |
| Bromeliad1776 | 72 | 209,186 | 6601.7 | 170,893<br>(81.7%) | 35,790<br>(17.1%) | 2,503<br>(1.2%) | 162,924<br>(77.9)% | 37,661<br>(18.0%) | 8,601<br>(4.11%) |
| <u>pop-level vcf</u> |  |  |  |  |  |  |  |  |  |
| Angiosperms353 | 38 | 15,622 | 1,837.8 | 13,345<br>(85.5%) | 1,442<br>(9.2%) | 835<br>(5.3%) | 3,032<br>(19.4%) | 1,129<br>(7.22%) | 11,461<br>(73.4%) |
| Bromeliad1776 | 40 | 65,473 | 3914.9 | 54,636<br>(83.5%) | 9,967<br>(15.2%) | 870<br>(1.3%) | 51,405<br>(78.5%) | 10,588<br>(16.2%) | 3,480<br>(5.3%) |

### 3.4 Similar phylogenomic resolution in concatenation method, Bromeliad1776 outperforms Angiosperms353 for species tree reconstruction

The Angiosperms353 and Bromeliad1776-based maximum-likelihood phylogenetic trees recovered the same backbone phylogeny of Bromeliaceae, clustering subfamily Tillandsiaoedeae and the subgenus *Tillandsia* with high bootstrap values (Supporting information Figure S5). Neither set obtained high support for inter-population structure for *Tillandsia gymnobotrya*, but highly-supported nodes separated *T. fasciculata* accessions from Mexico and from other locations, and the populations of *T. punctulata* for the Bromeliad1776 data set were similarly separated. The tree topologies were identical, with the notable exception of the placements of*Tillandsia biflora* and *Racinaea ropalocarpa* and the genus *Deuterocohnia* (Supporting information Figure S5, purple arrow). Overall, internal nodes are strongly supported for both sets, except for *Hechtia carlsoniae* as sister to Tillandsioideae, which is poorly supported for both sets. While several internal nodes are slightly less supported for the Angiosperms353 set, overall these results demonstrate the efficacy of both kits in phylogenomic reconstruction using concatenation approaches, indicating that as few as 47k SNPs within variable regions provide reliable information to resolve phylogenetic relationships within the recent evolutionary radiation of *Tillandsia*.

Species trees as inferred with ASTRAL for both data sets likewise provided an overall strong local posterior support (Figure 2, see also Supporting information). Several nodes however exhibit lower local posterior support values for the Angiosperms353 tree than for the Bromeliad1776 tree. The topology for the Bromeliad1776 ASTRAL tree was similar to the ML tree, but differed again by placing *Deuterocohnia* as sister taxa to *Puyoideae* only. In the Angiosperms353 tree, the topology differed from both ML trees and the ASTRAL Bromeliad1776 tree in several nodes. *H. carlsoniae* was placed as a sister taxa to all other subfamilies in the Angiosperm353 phylogeny. Notably, the placement of *Catopsis* and *Glomeropitcrania* differed, as well as the placement of *Cipurosis subandinai*, *T. biflora* and *R. ropalocarpa*. Several internal nodes were poorly supported, such as the node separating the tribe Catopsideae and core Tillandsioideae, and the nodes separating Tillandsioideae from all other subfamilies. The differences in topology between the Angiosperms353 ASTRAL tree to all other trees (ML trees and Bromeliad1776 ASTRAL tree) together with the low posterior support suggest lower resolution power and a poor fit of this data set for resolving a species tree.

**Figure 2.**
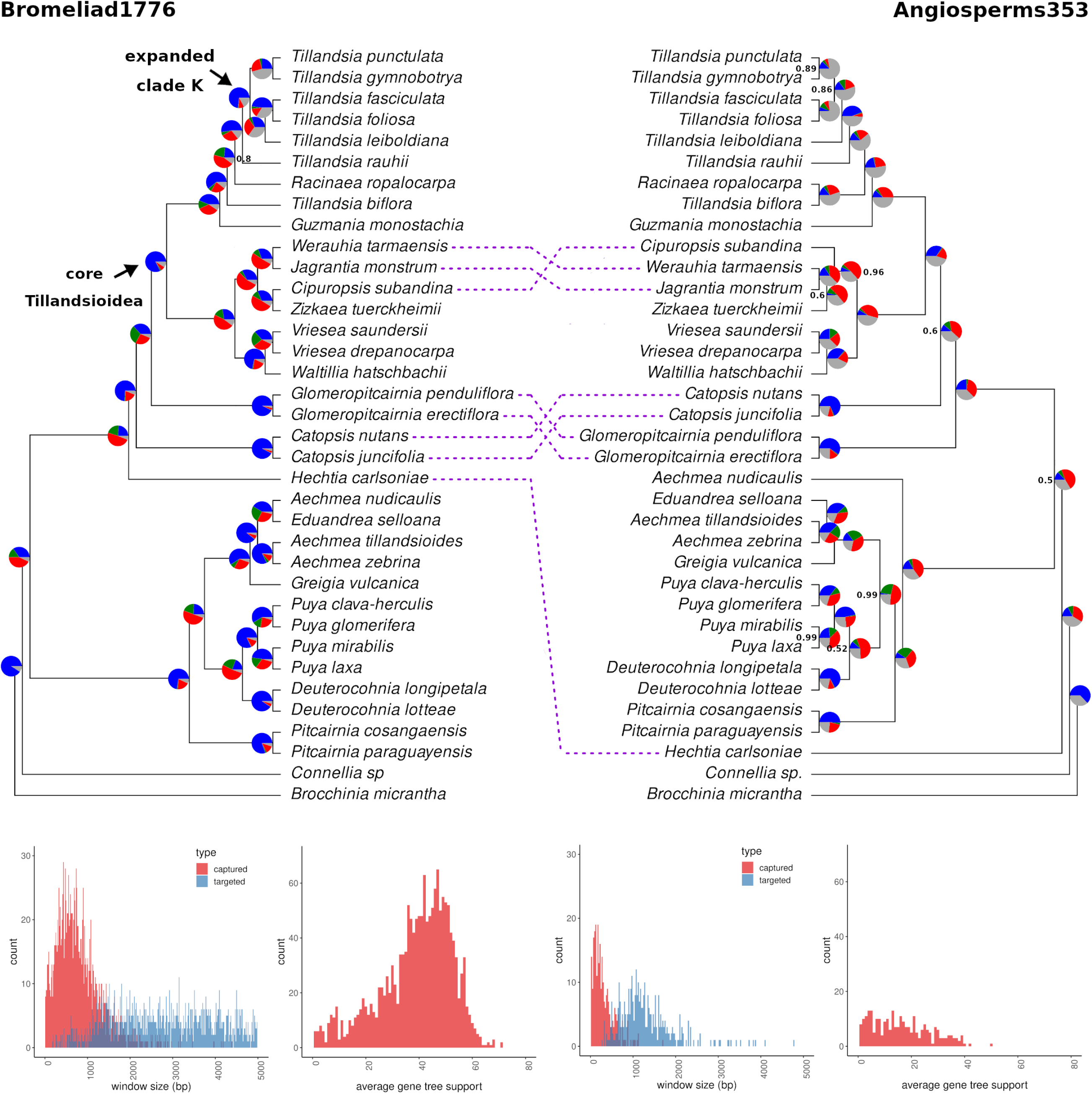
Coalescent-based species trees generated ASTRAL-III for samples enriched with Bromeliad1776 (left) and Angiosperms353 (right, flipped for mirroring), on 269 and 1600 genes for each set, respectively. Node values represent local posterior probabilities (pp) for the main topology and are equal to 1 unless noted otherwise. Pie charts at the nodes show levels of gene tree discordance: the percentages of concordant gene trees (blue), the top alternative bipartition (green), other conflicting topologies (red) and uninformative gene trees (gray). At bottom, length and average bootstrap support for gene trees from either data set, according to the design of the bait set used for enrichment: Angiosperms353 (right) and Bromeliad1776 (left). Each gene was considered a single genomic window.

The length and average size of the input gene trees different among sets, with average window length of 304.6 bp and 819.9 bp and average gene tree support of 16.9 and 38.9 for Angiosperms353 and Bromeliad1776 bait-sets, respectively (Figure 2). An examination of gene tree concordance constructed with Bromeliad1776 data set allowed us to identify variable levels of gene tree conflict among nodes (Figure 2). Gene tree discordance was especially high for the split between Tillandsioideae and other subfamilies, as well as for the split between Puyoideae and taxa assigned to Bromelioideae. Furthermore, gene tree discordance and the proportion of un-informative gene trees was especially high for splits among clades within the K.1 and K.2 clades of subgenus *Tillandsia*. A similar analysis with Angiosperms353 yielded evidence for gene tree discordance, but a considerable number of gene trees were reported to be non-informative (grey part of the pie charts), especially within subgenus *Tillandsia* (Figure 2).

### 3.5 Strong interspecific structure, but little evidence for within-species population structure

After LD-pruning and retaining maximum 10% missing data, 1,025 and 32,941 biallelic SNPs were included for the *Tillandsia* PCA analysis of the Angiosperms353 and Bromeliad1776 data sets, respectively. Overall, both data sets provided evidence for interspecific structure, but not for population structure, with Bromeliad1776 resulting in border-line higher resolution (slightly better separating *T. foliosa* from *T. fasciculata*). The percentage of explained variance was higher in the Bromeliad1776 set (19.3% and 16.5% for PC1 and PC2) as compared to the Angiosperms353 data set (14.5% and 11.8%, see Figure 3, Supporting information Figure S6). Based on these two PCAs, we found no evidence for spatial genetic structure within each species, since accessions did not cluster by geographic origin on the two PCs presented, or any other PCs we investigated (See Supporting information Figure S6).

**Figure 3.**
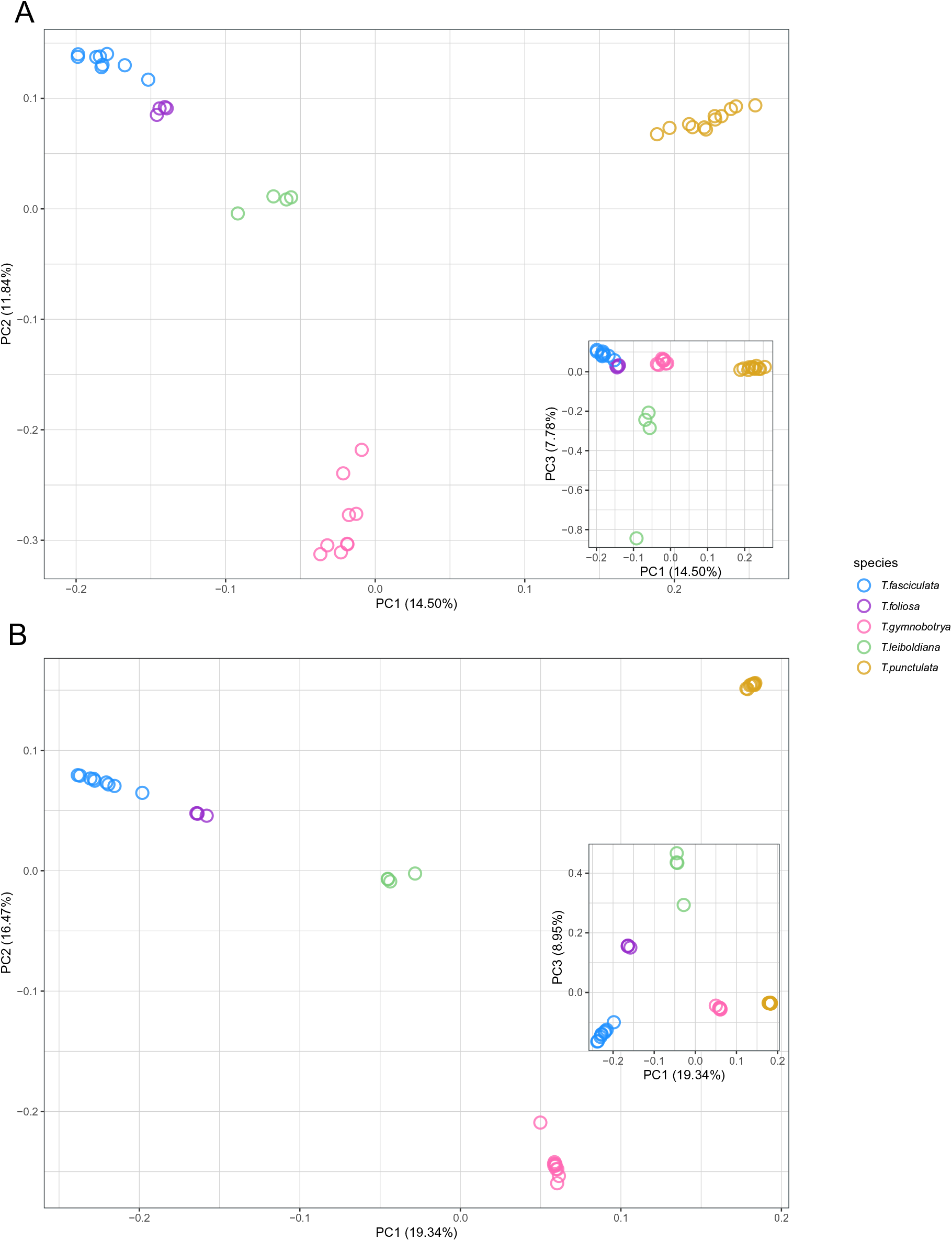
Principal Component Analysis (PCA) plot for samples of Tillandsia subgenus Tillandsia enriched with two bait sets: A. Angiosperms353 (1,025 variants); B. Bromeliad1776 (32,941 variants). Colors indicate different species according to legend.

In addition to PCA, we performed ADMIXTURE analyses based on 9,804 and 42,613 variants for the Angiosperms353 and Bromeliad1776 sets, respectively (Figure 4). We used a cross-validation strategy to identify the best K and found clear support for K=5 for the Bromeliad1776 set (Supporting information Figure S7). In contrast, the CV pattern for the Angiosperms353 set varied widely, providing limited information about the best K. Lowest CV values were however observed for K=9 with locally low values for K=5 and K=3 (Supporting information Figure S7). We further investigated the ADMIXTURE bar plots at different values of K. For K=5, very similar patterns can be observed for both sets, with the recovered clusters reflecting the expected species boundaries. The main difference between the two data sets was the ability of the Bromeliad1776 set to reach a more consistent solution (“consensus”) among 30 runs, especially at large K, as compared to the runs based on the Angiosperms353 bait set. The Bromeliad1776 was also able to distinguish between different sampling localities of *T. punctulata* and of *T. fasciculata* at K=7-8 (Figure 4).

**Figure 4.**
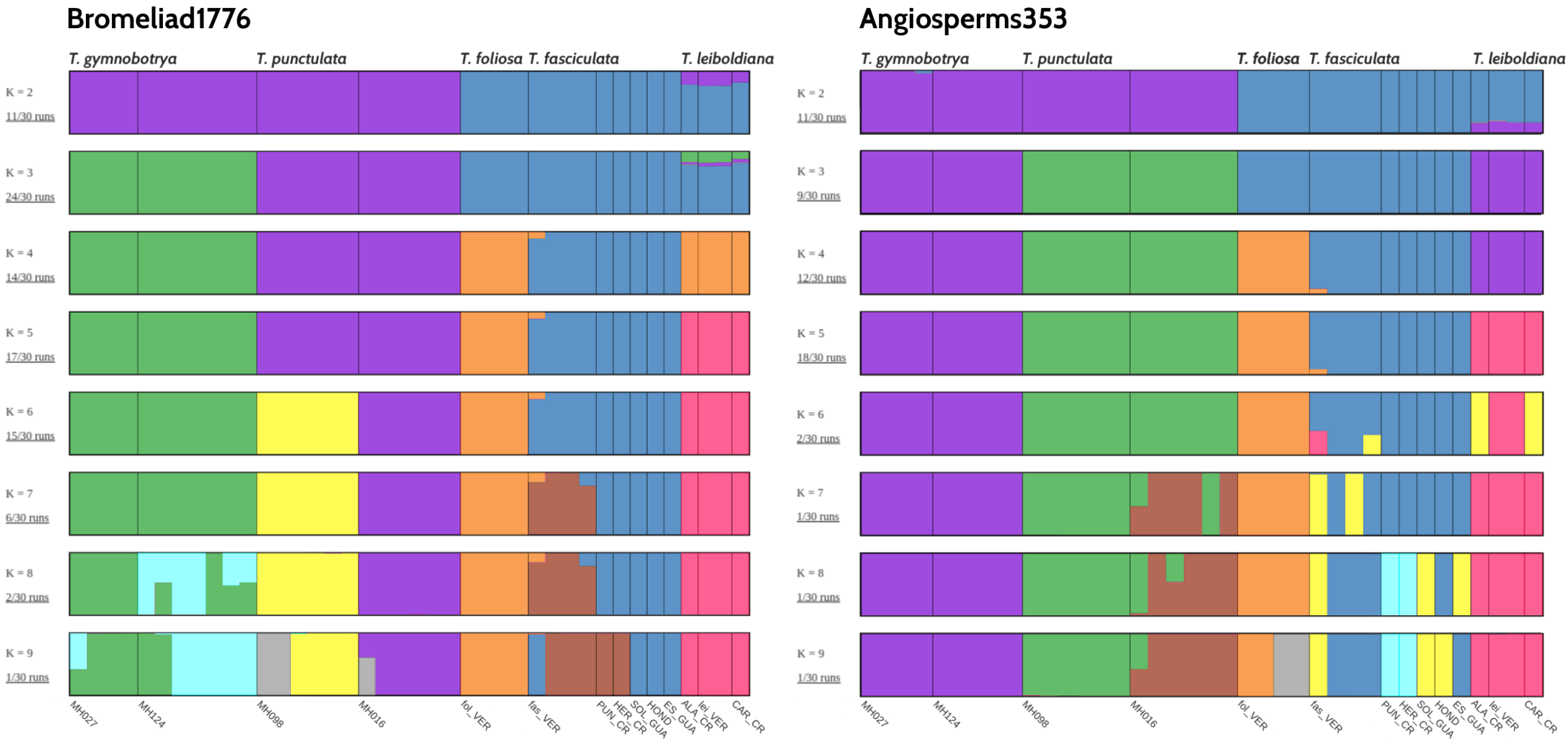
Population structure of 5 Tillandsia subgenus Tillandsia species from 14 sampling locations inferred with the ADMIXTURE software. Samples were enriched with either of two bait sets: Angiosperms353 (9,804 variants after LD-pruning) and Bromeliad1776 (42,613 variants after LD-pruning), showing values of K=2 to K=9. Colors represent genetically differentiated groups while each accession is represented by a vertical bar.

### 3.6 Distinct diversities hint at different demographic processes

Nucleotide diversity estimates were calculated for the Bromeliad1776 data-set only, due to difficulties obtaining a reliable SNP set with Angiosperms353 (see section 2.4). Averaged levels of nucleotide diversity at synonymous sites*π*_S_ greatly varied among species, from 4.1*x*10*^−^*^3^ to 8.1*x*10*^−^*^3^ for *T. foliosa* and *T. fasciculata*, respectively (Supporting information Table S5; Figure 5). Given the recent divergence of these different species and their roughly similar life history traits, they are expected to share relatively similar mutation rates, hence the observed differences in *π*_S_ are expected to translate into differences of long-term N_e_. Looking at the distribution of *π*_S_ across genes, we foundbroader or narrower distributions depending on the species, which explains the observed differences in averaged *π*_S_, as typically represented by the median of the distribution (vertical bars, Figure 5). Most species exhibit distributions of Tajima’s D (Fig 5) that are centered around zero, with the notable exception of *T. punctulata*. The distribution of this species is shifted toward positive Tajima’s D values, therefore indicating a recent population contraction, suggesting that this species experienced a unique demographic trajectory as compared to the other species.

**Figure 5.**
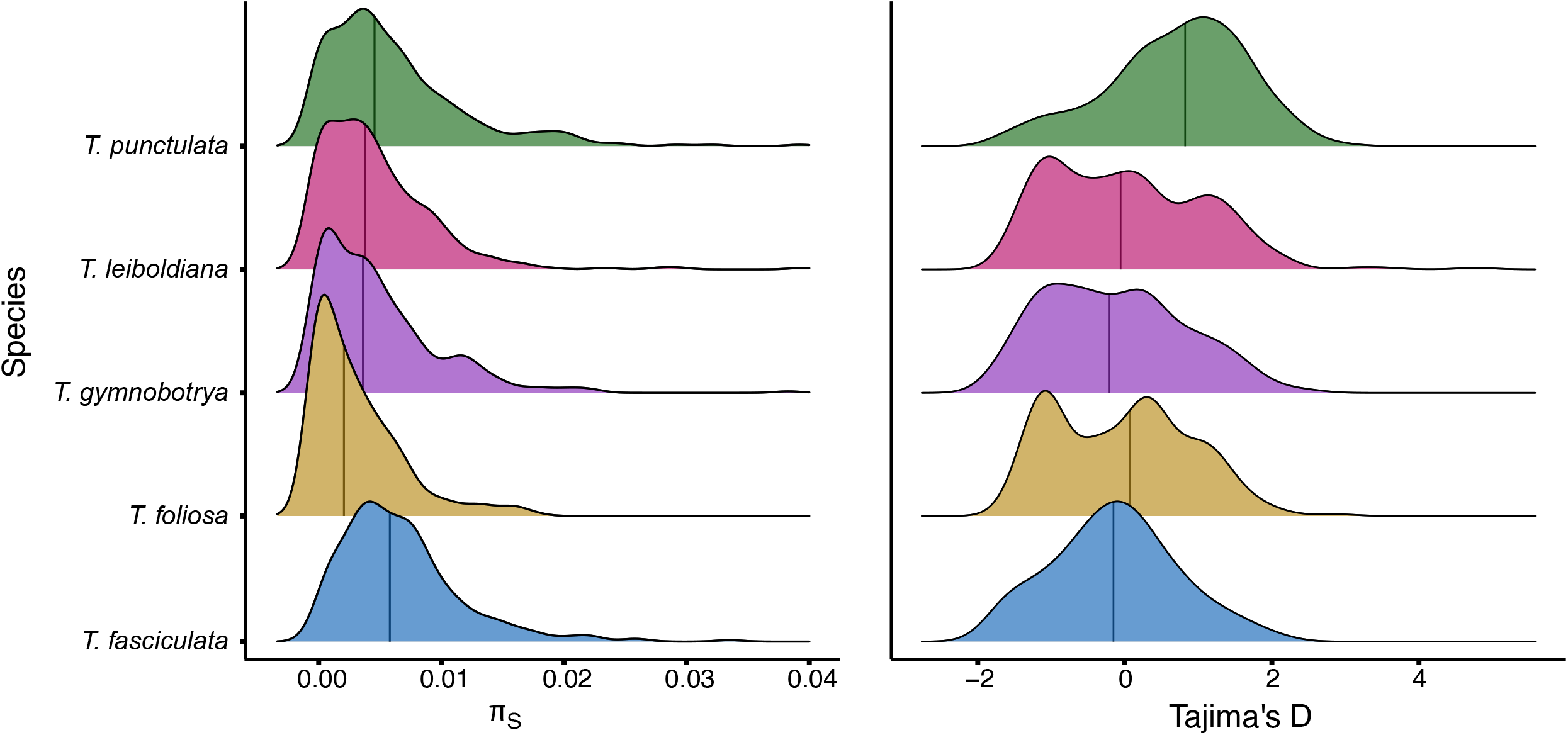
Distribution of Tajima’s D and synonymous (π_S_) nucleotide diversity within each species for the Bromeliad1776 kit.

## 4 Discussion

### 4.1 A taxon-specific bait set performs marginally better for phylogenomics

In this study, we compared the information content and performance of a taxon-specific bait set and a universal bait set for addressing questions on evolutionary processes at different scales in a highly diverse Neotropical plant group, including recently radiated clades. We found that the taxon-specific kit provided a greater number of segregating sites, yet contrary to our expectations, the abundance of information content did directly translate to a greater resolution power.

The universal and taxon-specific sets performed comparably when investigating macroevolutionary patterns: the inferred species trees are remarkably consistent between the two bait sets (Supporting information Figure S5, Figure 2). Notably, both sets were sufficiently informative to reconstruct the relationships among the fastest radiating clades. These results resonate with previous comparative works (e.g. in *Burmeistera*, Bagley et al., 2020; in *Buddleja,* Chau, Rahfeldt, & Olmstead, 2018; and in *Cyperus*, Larridon et al., 2020), where taxon-specific markers provided higher gene assembly success, but a comparable number of segregating sites for phylogenetic inference, indicating that universal bait sets are nearly as effective as taxon-specific bait sets, even in fast evolving taxa. The main advantage of the bromeliad taxon-specific set is its ability to provide additional resolution for deeper examination of gene tree incongruence (Figure 2), currently a fundamental tool in phylogenomic research (Edwards, 2009; Morales-Briones et al., 2020; Pease et al., 2016).

The taxon-specific bait set performed marginally better to address hypotheses at more recent evolutionary scales and provided arguably clearer evidence for inference of species genomic structure using clustering methods. In fact, genetic markers obtained from both data sets provided sufficient information to infer species but no geographic structure, suggesting that *Tillandsia* could be characterized by high gene dispersal among populations. Considering that the Angiosperms353 kit has shown potential to provide within-species signal, as recently demonstrated by Beck et al. (2021) on *Solidago ulmifolia*, and to estimate demographic parameters from herbarium specimen (Slimp et al., 2021), we would expect the taxon-specific set to accurately reveal a geographical genetic structure. However, the present study is generally based on small sample sizes per species (n=4-8), mostly sampled within a limited geographic range, limiting our ability to draw robust conclusions on the levels of intra-specific population structure.

The Bromeliad1776 kit provided a substantially larger number of segregating sites (more than 200k vs. 47k in Angiosperms353; Table 1, Supporting information Figure S2) due to higher enrichment success, following the expectation for higher sequence variation in custom-made loci (Figure 1, see also Bragg et al., 2016; de La Harpe et al., 2019; Kadlec et al., 2017). We accordingly found that rates of molecular divergence are distinctly correlated with enrichment success in our sampling (Figure 1), following the expectation that a universal kit will provide fewer segregating sites.

However, the difference in resolution power between the kits cannot be ascribed solely to the different numbers of SNPs, but rather to the length and variability of the obtained regions. The topology obtained with the Angiosperm353 data set under the multi-species coalescent model was substantially different from all other inferred trees and the input gene trees provided a low power to detect patterns of gene tree discordance (Figure 2). We additionally observed that the highly conserved regions targeted by Angiosperms353 are shorter in comparison to Bromeliad1776 targets and thus result in shorter input windows for species tree inference (Figure 2). Hence, the patterns of gene tree discordance in the Angiosperms353 data set likely indicate incorrect gene tree estimation or other model misspecifications, rather than a biological signal. Specifically, coalescence-based methods are sensitive to gene tree estimation error (Zhang et al., 2018) and perform better with gene trees estimated from unlinked loci long enough and variable enough to render sufficient signal per gene tree - this is especially true for data sets with many taxa. The high rates of uninformative genes trees, found in almost half of the intergenic nodes in the Angiosperms353 data set, is expected with increased levels of gene tree error which in turn reduce the accuracy of ASTRAL (Mirarab, 2019; Sayyari & Mirarab, 2016). In contrast, the Bromeliad1776 ASTRAL tree (Figure 2, left) resolved phylogenetic relationships among taxa with high posterior probability and a topology similar to the ML tree. Gene tree discordance analysis revealed high incongruence around certain nodes, possibly reflecting rapid speciation events.

Since inference of phylogenetic relationships under the multi-species coalescent and exploration of gene tree discordance are both pivotal to phylogenomic research (Degnan & Rosenberg, 2009; Edwards et al., 2016; Pease et al., 2016), a taxon-specific kit provides a clear advantage especially in recent rapid radiations, where gene tree conflict and incomplete lineage sorting are expected to be prevalent (Dornburg, Su, & Townsend, 2019; Kubatko & Degnan, 2007; Roch & Warnow, 2015). In that regard, inference of the species tree with the Bromeliad1776 is a tool to drive further hypotheses concerning evolutionary and demographic processes in the evolution of *Tillandsia*. Moreover, the features of the loci targeted provide an important opportunity to study selection (see section 4.3).

### 4.2 Insights on Bromeliaceae phylogeny and demographic processes in *Tillandsia*

Both bait sets resolved the phylogeny of Bromeliaceae, including the fastest evolving lineages of the subfamily Tillandsioideae. The results generally agreed with previous findings of the relationships among taxa (Givnish et al., 2011, 2014). Several findings that contrast with the expected known phylogeny may point at a complexity of genomic processes in the evolutionary history of Bromeliaceae subfamilies. Both the ML tree and species tree did not support a monophyly of the subfamily Pitcairnioideae, which was represented by four samples and two genera in our phylogeny: *Deuterochonia* and *Pitcarnia*. Rather, the genus *Deuterochonia* was sister to subfamily Puyoideae or sister to both Puyoideae and Bromelioideae subfamilies, inconsistent with the results of Barfuss et al. (2016) and Granados Mendoza et al. (2017). Interestingly, in a visualization of gene tree discordance we found high levels of incongruence and a high percentage of trees supporting an alternative topology in the node splitting the genera, indicating that several genomic processes such as hybridization and incomplete lineage sorting may have accompanied divergence in this group, contributing to the phylogenetic conflict and extending the challenges in resolving these evolutionary relationships. Within the core Tillandsioideae, the tribes Tillandsieae and Vrieseeae were found to be monophyletic, in accordance with previous work on the subfamily (Barfuss et al., 2016). Finally, within our focal group *Tillandsia* subgenus *Tillandsia*, clade K as suggested by Barfuss et al. (2016) and clades K.1 and K.2 as proposed by Granados Mendoza et al. (2017) were all well supported, further in agreement with their interpretation of Mexico and Central America as a center of diversity for subgenus *Tillandsia*. Within *Tillandsia*, incongruence was prominent at the recent splits within clade K.1. and clade K.2 as expected in a recent rapid radiation, a result of high levels of incomplete lineage sorting, hybridization and introgression (Berner & Salzburger, 2015).

When applied to methods in population genetics, we obtained some evidence for a difference in demographic processes and in the level of genetic variation among species. This was especially true for the taxon-specific bait set: for example, the bait set differentiated between populations of *T. punctulata* and *T. fasciculata*, but not *T. gymnobotrya* in a maximum likelihood tree and ancestry analysis (Supporting information Figure S5, Figure 4), indicating differences in inter-population genetic structure among species. The evidence for different demographic processes in these species extended to estimates of Tajima’s D, where lower values may indicate a recent bottleneck. In addition, we found a unique distribution of nucleotide diversity for *T. foliosa*, possibly reflecting a low effective population size for this endemic species in contrast with the closely related, but widespread *T. fasciculata*. In all cases, our limited sampling given the large size of the family constrains our ability to draw conclusions of a ‘true’ phylogeny and to account for population structure. Our finding however suggests that nuclear markers obtained with a target capture technique can highlight genomic processes and be further applied to address questions in population genomics with a wider sampling scheme.

### 4.3 Future prospects and implications for research in Bromeliaceae and rapid radiations

Beyond the scope of this study, the availability of a bait set kit for Bromeliaceae provides a prime genetic resource for investigating several topical research questions on the origin and maintenance of Bromeliaceae diversity. Manyfold studies of bromeliad phylogenomics set forth the challenges of resolving species-level phylogenies with a small number of markers, particularly in young and speciose groups (Goetze, Zanella, Palma-Silva, Büttow, & Bered, 2017; Granados Mendoza et al., 2017; Loiseau et al., 2021; Versieux et al., 2012). This particularly curated bait set allows highly efficient sequencing across taxa: within our study, we found high mapping success with 82.3% average read mapping. As expected, we documented a difference in enrichment success among taxa, explained by divergence time to the reference used for bait design (see Supporting information Figure S4), suggesting possible deviations from the assumptions of non-randomly distributed missing data that may mislead phylogenetic inference (A. R. Lemmon, Brown, Stanger-Hall, & Lemmon, 2009; Streicher, Schulte, & Wiens, 2016; Xi, Liu, & Davis, 2016). However, given the large enrichment success, downstream analysis with deliberate methodology can account for possible biases and provide robust inference with strict data filtering (Molloy & Warnow, 2018; Streicher et al., 2016). Hence, target enrichment with Bromeliad1776 can produce large data sets with consistent representation between taxa, allowing repeatability between studies and retaining the possibility for global synthesis by including sequence baits orthologous to the universal Angiosperms353 bait set. Moreover, with specific knowledge of the loci targeted in this set, the ability to obtain the same sequences across taxa and experiments and to differentiate genic regions with the use of *A. comosus* models, this bait set offers a broad utility for research in population genomics.

Another important feature in the Bromeliad1776 set is the inclusion of genes putatively associated with key innovative traits in Bromeliaceae with a focus on C3/CAM shifts. Little is known about the molecular basis of the CAM pathway, an adaptation to arid environments which evolved independently and repeatedly in over 36 plant families (Heyduk, Moreno-Villena, Gilman, Christin, & Edwards, 2019; Chen, Xin, Wai, Liu, & Ming, 2020; Silvera et al., 2010). CAM phenotypes are considered key adaptations in Bromeliaceae, associated with expansion into novel ecological niches. In *Tillandsia*, C3/CAM shifts were found to be particularly associated with increased rates of diversification (Crayn et al., 2004; de La Harpe et al., 2020; Givnish et al., 2014). The Bromeliad1776 bait set offers opportunities to address specific questions on the relationship between rapid diversification and photosynthetic syndromes in this clade, including testing for gene sequence evolution. Additionally, the inclusion of multi-copy genes, combined with newly developed pipelines for studying gene duplication and ploidy (Morales-Briones et al., 2020; Viruel et al., 2019), are beneficial for studying the role of gene duplication and loss in driving diversification. With the increasing ubiquity of target baits as a genomic tool we expect to see additional pipelines and applications emerging, further expanding the utility of target capture for both macro-and microevolutionary research.

## 5 Conclusions

Even as whole genome sequencing becomes increasingly economically feasible, target capture is expected to remain popular due to its extensive applications in research. We found that evaluating the differences in resolution power between universal and taxon-specific bait sets is far from a trivial task, and we attempted to lay out a methodological roadmap for researchers wishing to reconstruct the complex evolutionary history of rapidly diversifying lineages. While a taxon-specific set offers exciting opportunities beyond phylogenomic and into research of molecular evolution, its development is highly time-consuming, requires community-based knowledge and may cost months of work when compared with out-of-the-box universal kits. Our results suggest that universal kits can continue to be employed when aiming to reconstruct phylogenies, in particular as this may offer the possibility to use previously published data to generate larger data sets. However, for those wishing to deeply investigate evolutionary questions in certain lineages, a taxon-specific kit offers certain benefits during data processing stages, where knowledge of the design scheme and gene models is extremely useful, and the possible return of costs is especially high for taxa emerging as model groups. We furthermore encourage groups designing taxon-specific kits to include also universal probes, furthering the mission to complete the tree of life.

## 6 Acknowledgments

This paper is dedicated to Christian Lexer, a wonderful mentor, friend and colleague. The research was supported with funding from the Christian Lexer professorship startup BE772002. The analyses benefited from the Vienna Scientific Cluster (VSC) and the Montpellier Bioinformatics Biodiversity (MBB) platform services. We thank Huiying Shang and Aram Drevekenin for help with code, Kelly Swarts, Claus Vogl and Matt Johnson for insightful discussions and advice. We thank the members of Swiss SNSF Sinergia project CRSII3_147630 for accession sampling and three anonymous reviewers for their comments on an earlier version of this manuscript.

## 7 Data Accessibility

Targeted sequencing reads generated for this project are available at NCBI-SRA under Bio-Project PRJNA759878; for accession numbers, see supporting information Table S4. The probe set and the relevant supporting information are available in Dryad (doi:10.5061/dryad.mpg4f4r11). The bioinformatics scripts are available at https://github.com/giyany/Bromeliad1776/tree/main/MS_2021_scripts.

## 8 Author Contribution

CL, MP and GY conceived the study. CL provided funding. TK coordinated sample collection, MdLH, VGJ and GY collected data. Species identified by MHJB and WT. Bait kit designed by GY, with guidance from JH and MP. Molecular work was performed by CGC, JV, NR, MHJB and GY. The data was analyzed by GY and TL with feedback from JV and OP. The manuscript was written by GY with significant input from all co-authors.

## 10 Supporting information

### 10.1 Tables

**Table S1** Genes included in the Bromeliad1776 bait design, with identifiers as annotated in *Ananas comosus* genome v.3 (Ming et al., 2015). The table includes details about exon composition, copy number and putatively associated pathways.

**Table S2** Categories of pathways and traits used to choose genes of interest for the Bromeliad1776 bait set, including literature source and number of genes in each category.

**Table S3** List of accessions used in this study. For samples of Tillandsia subgenus Tillandsia locality codes are also indicated.

**Table S4** Number of reads, numbers and percentage of read mapping to target in all samples for both bait sets.

**Table S5** Averaged levels of nucleotide diversity at synonymous (*π*_S_) and non-synonymous (*π*_N_) for 5 *Tillandsia* subgenus *Tillandsia* species.

### 10.2 Figures

**Figure S1** Map of sampling locations for *Tillandsia* subgenus *Tillandsia* accessions within Mexico.

**Figure S2** Mapping rates (A) and percentage of reads matching bait sequences (B) for Bromeliad samples enriched with one of two bait sets: Angiosperms353 and Bromeliad1776. Reads were mapped against *A. comosus* reference for both bait sets. Targets were defined as bait locations and flanking 500 base-pairs. Bromeliad1776 targets were defined as the regions used for bait design and Angiosperms353 targets were defined as *A. comosus* orthologous regions matching the genes used for bait design.

**Figure S3** A simplified phylogenetic tree, with branches colored according to read mapping percentage for samples enriched with Angiosperms353.

**Figure S4** A simplified phylogenetic tree, with branches colored according to read mapping percentage for samples enriched with Bromeliad1776.

**Figure S5** Maximum-likelihood (ML) phylogenetic tree inferred with RAxML-NG, based on variants called for data sets enriched with Bromeliad1776 bait set (left) and Angiosperms353 bait set (right, flipped for mirroring). Branch lengths were calculated by number of substitutions per site. Internal nodes are marked and colored according to bootstrap support. Nodes which differed among trees are colored purple and have been marked by an arrow.

**Figure S6** Principal Component Analysis (PCA) plot for samples of *Tillandsia* subgenus *Tillandsia* enriched with two bait sets: A. Angiosperms353 (1,025 variants after LD-pruning) B. Bromeliad1776 (32,941 variants after LD-pruning). Colors indicate different species (following the scheme in Supporting Figure S6) and shapes represent different geographic origins (populations).

**Figure S7** Admixture cross-validation errors (top) detected for values of K between 2 and 9 for A. Angiosperms353 data set and B. Bromeliad1776 data set.

**Figure S8** Coalescent-based species trees generated ASTRAL-III for samples enriched with Angiosperms353 using 269 genes. Node values represent local posterior probabilities (pp) for the main topology.

**Figure S9** Coalescent-based species trees generated ASTRAL-III for samples enriched with Bromeliad1776 using 1600 genes. Node values represent local posterior probabilities (pp) for the main topology.

### 10.3 Files

**File S1** Estimation of ascertainment bias in target capture data using comparison with whole-genome data

## Notes

### Competing Interest Statement

The authors have declared no competing interest.

### Summary of Updates

several clarifications on methodology used and background: changes in Fig 3.

