## Supplementary material for "Taxon-specific or universal? Using target capture to study the evolutionary history of rapid radiations": supp_info_ascertainment bias.docx

The non-random selection of markers for target capture may introduce ascertainment bias, with implications for population genomics methods (Heslot et al., 2013). An absence or misrepresentation of rare SNPs is an indication of such bias and may further result in loss of valuable information and misinterpretation of values, such as Fst and heterozygosity (Lachance & Tishkoff, 2013). To assess the extent of ascertainment bias that may rise in the use of Bromeliad1776, we assessed differences in SNP discovery by comparing whole genome data and target capture data in three *Tillandsia* samples, using the ratio of heterozygous to homozygous sites per LG as a measure to examine differences in SNP discovery.

Briefly, for each sample we obtained both whole-genome sequencing data and targeted sequencing data, produced with the bromeliad1776 bait-set. All accessions were reference mapped to the *A. comosus* v.3 reference (Ming, 2015). Variants were then called for each data type separately using freebayes v1.3.2-dirty (Garrison & Marth, 2012) - one variant call file for each data type, containing all three samples. Each vcf file was filtered to retain mean genotype depth > 4 and sites with 70% missing data using vcftools (Danecek et al., 2011). The ratio of heterozygous to homozygous sites was calculated for each individual and each LG (chr) using vcflib (Garrison, 2012).

The ratio heterozygous to homozygous sites was higher for target capture data sequencing in almost all species and loci, indicating high discovery of heterozygous sites. However, the differences in ratio were strongly correlated with sequencing depth (P < 0.003) and species (P < 0.003) and differences among species were more prominent than variation among data-types, indicating the former as causes for dissimilarities in SNP discovery, rather than marker selection (see figure below).

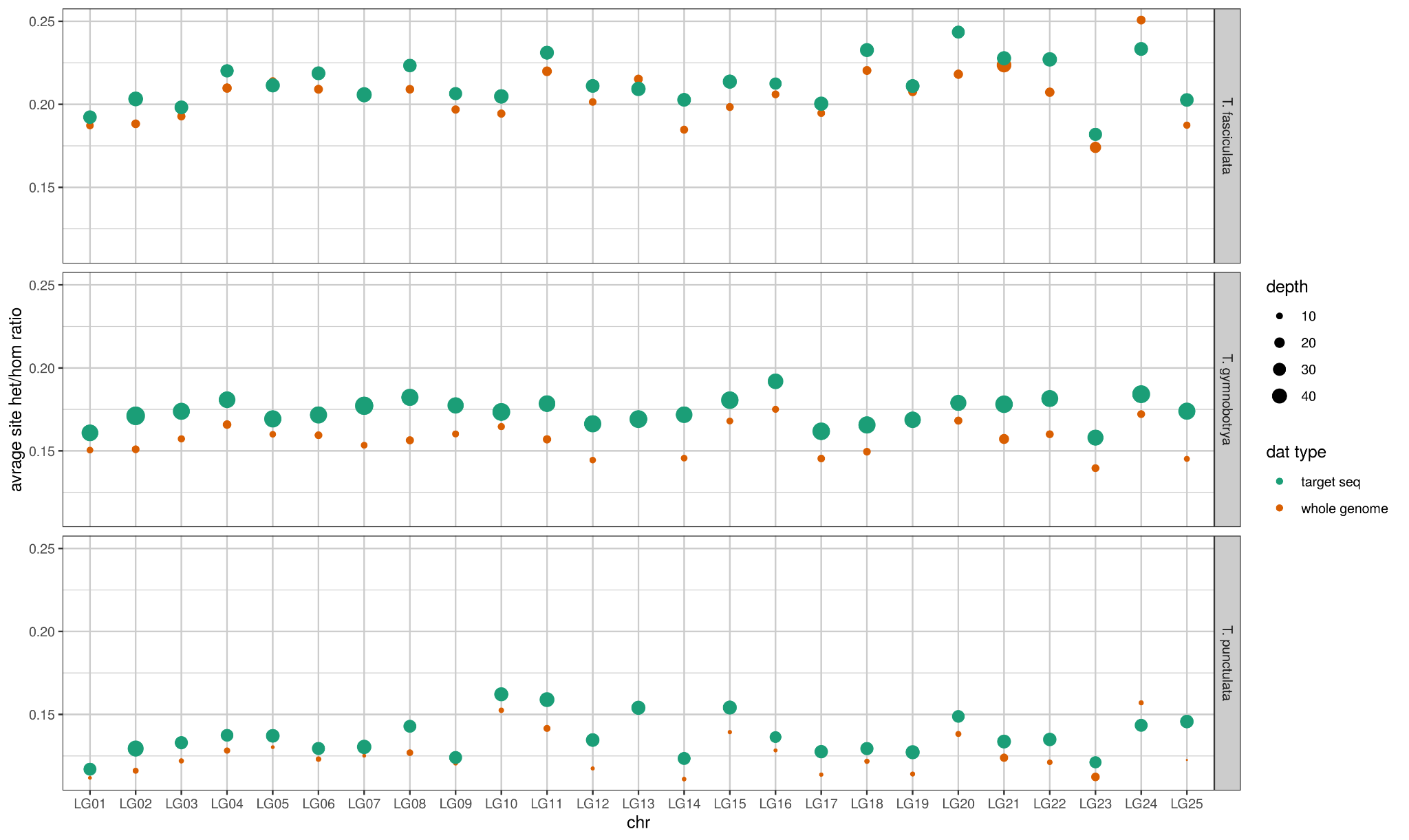

Danecek, P., Auton, A., Abecasis, G., Albers, C. A., Banks, E., DePristo, M. A., ... & 1000 Genomes Project Analysis Group. (2011). The variant call format and VCFtools. Bioinformatics, 27(15), 2156-2158.

Garrison, E. (2012). Vcflib: A C++ library for parsing and manipulating VCF files. GitHub https://github. com/ekg/vcflib.

Garrison, E., & Marth, G. (2012). Haplotype-based variant detection

from short-read sequencing. arXiv:1207.3907 [q-bio] .

Heslot, N., Rutkoski, J., Poland, J., Jannink, J.-L., & Sorrells, M. E.

(2013). Impact of Marker Ascertainment Bias on Genomic Selection

Accuracy and Estimates of Genetic Diversity. PLOS ONE , 8 (9),

e74612. doi: 10.1371/journal.pone.0074612

Lachance, J., & Tishkoff, S. A. (2013). SNP ascertainment bias in

population genetic analyses: Why it is important, and how to correct

it. BioEssays, 35 (9), 780–786. doi: 10.1002/bies.201300014

Ming, R., VanBuren, R., Wai, C. M., Tang, H., Schatz, M. C., Bowers,

J. E., . . . Yu, Q. (2015). The pineapple genome and the evolution

of CAM photosynthesis. Nature Genetics, 47 (12), 1435–1442. doi:

10.1038/ng.3435
