## Supplementary figures and images for "Taxon-specific or universal? Using target capture to study the evolutionary history of rapid radiations"

### Angio353_species_tree_t3_assigned_species.figtree.tre.pdf

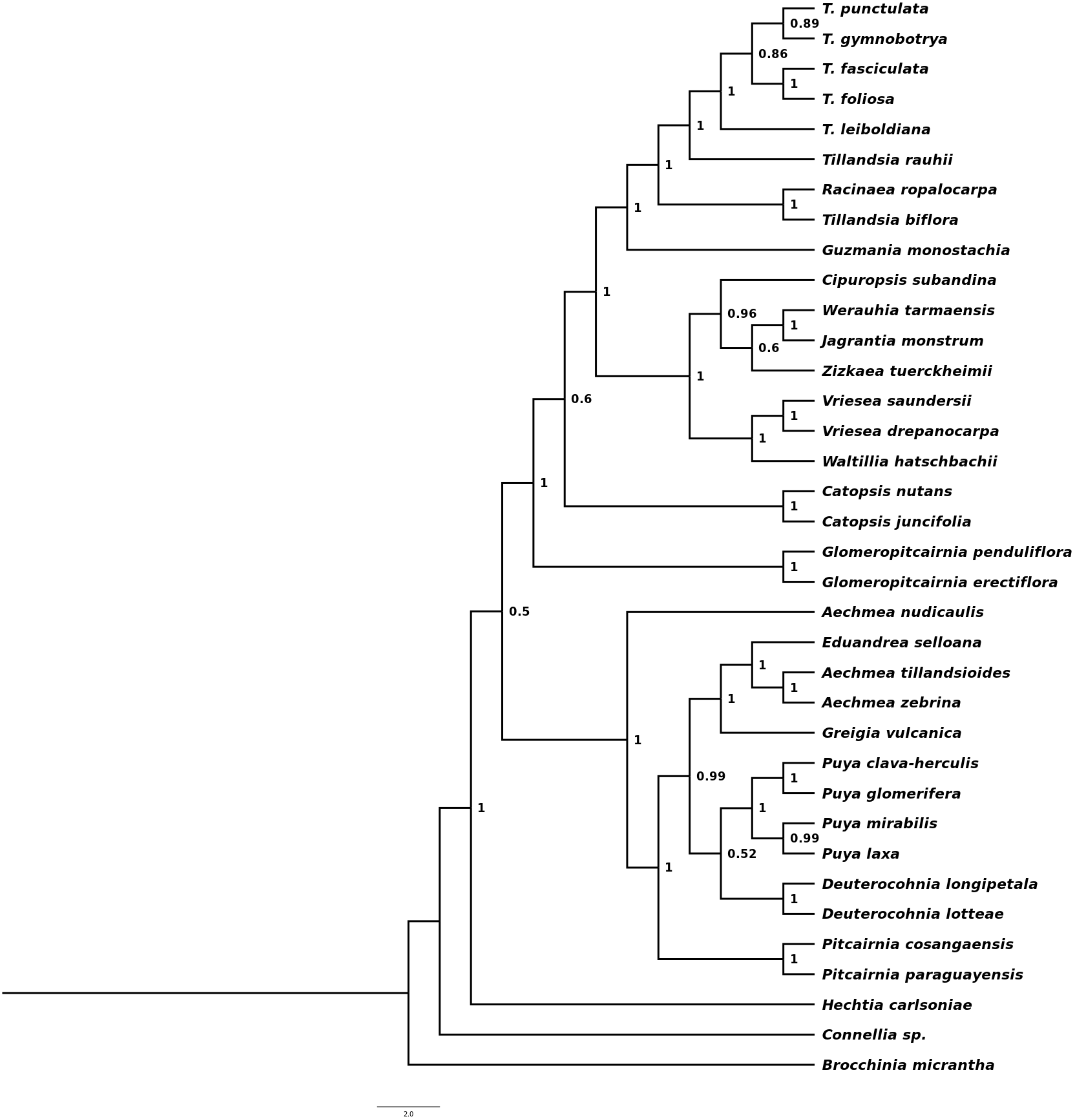

### Brom1776_species_tree_t3_assigned_species.figtree.tre.pdf

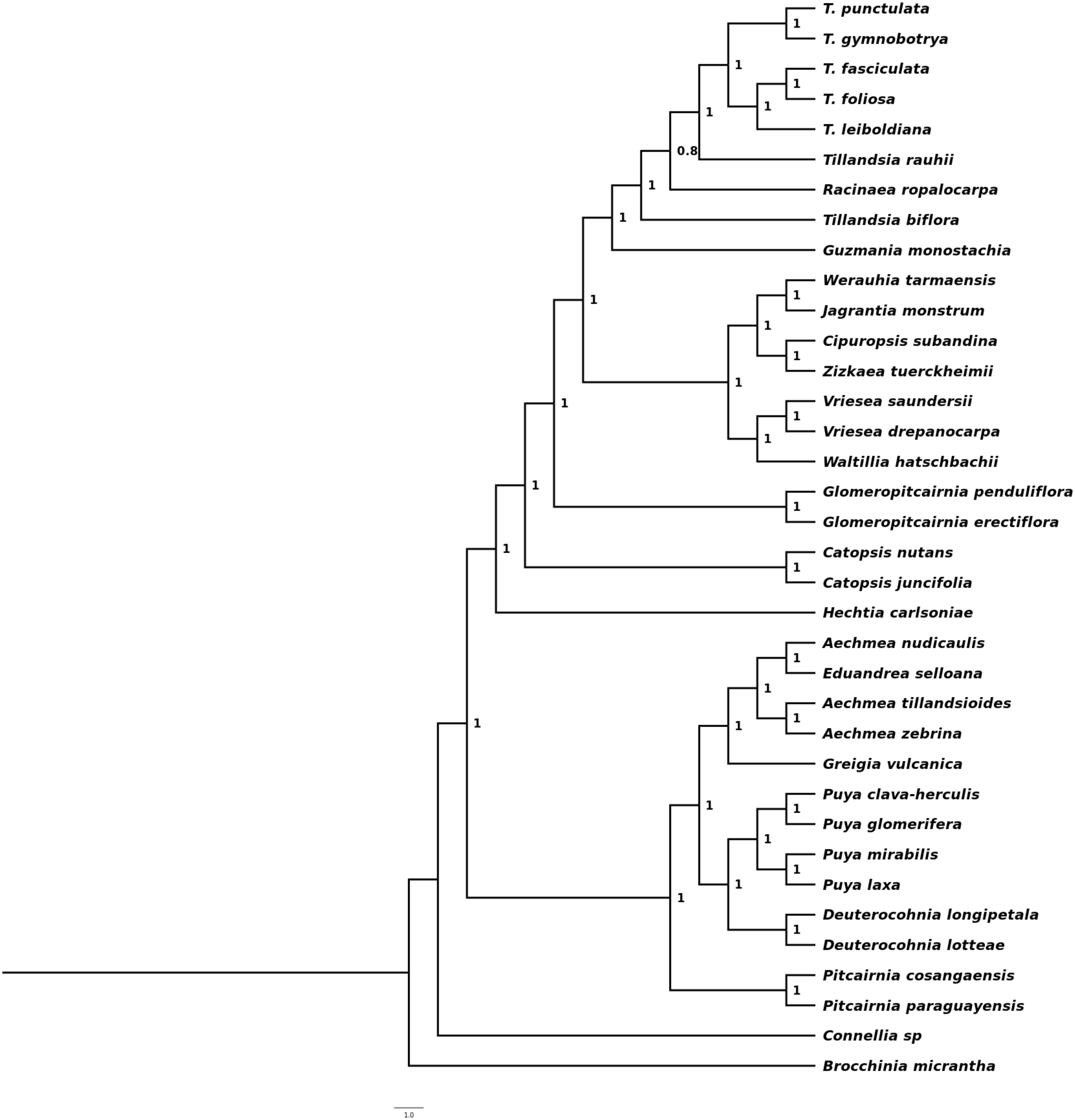

### supp_both_baitsets_phyparts_pies.pdf

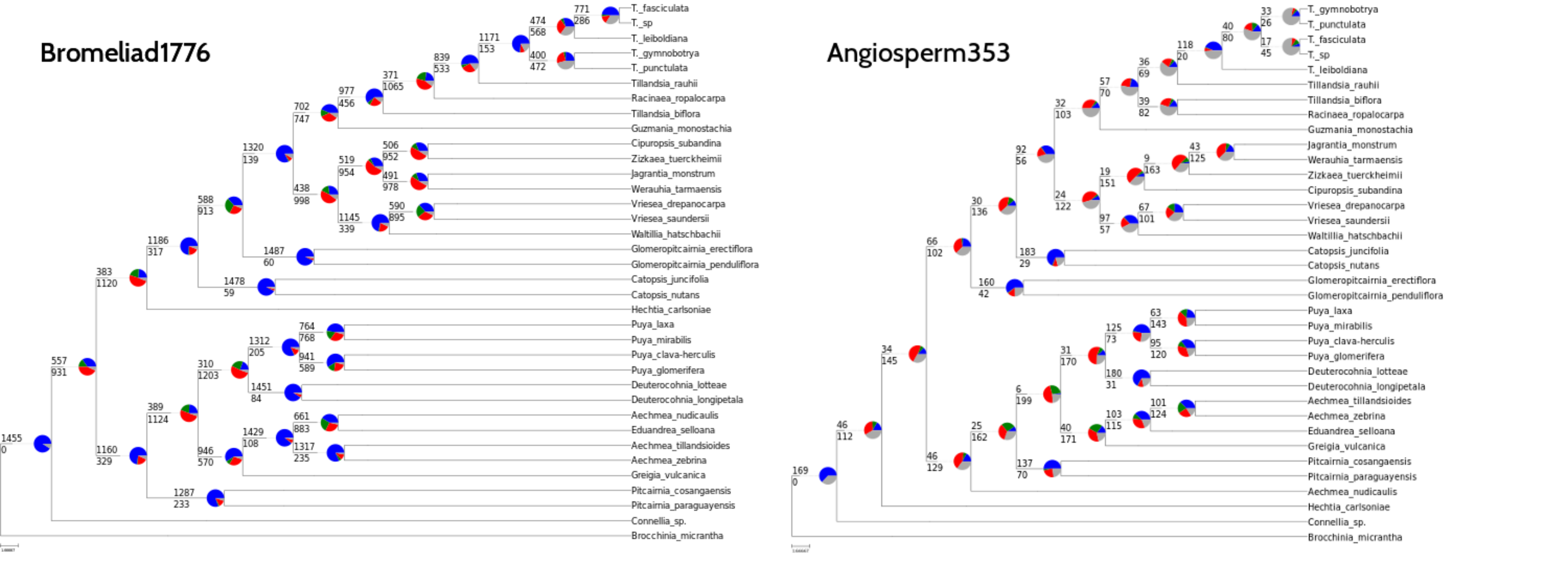
