## Supporting information - Figures 1-7 for "Taxon-specific or universal? Using target capture to study the evolutionary history of rapid radiations"

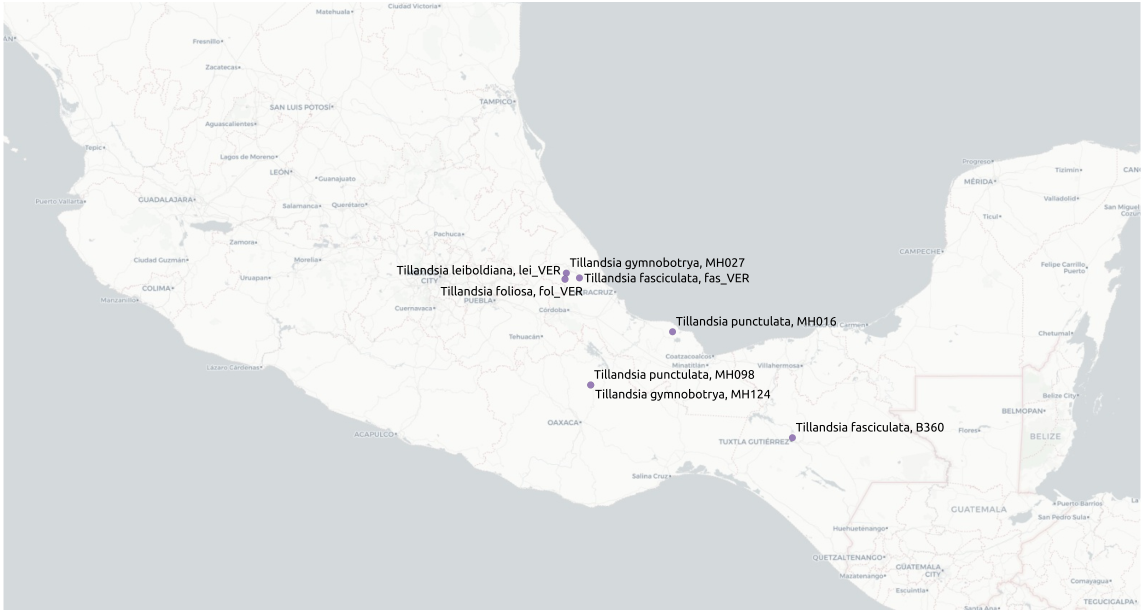


**Figure S1** Map of sampling locations for *Tillandsia* subgenus *Tillandsia* accessions within Mexico.

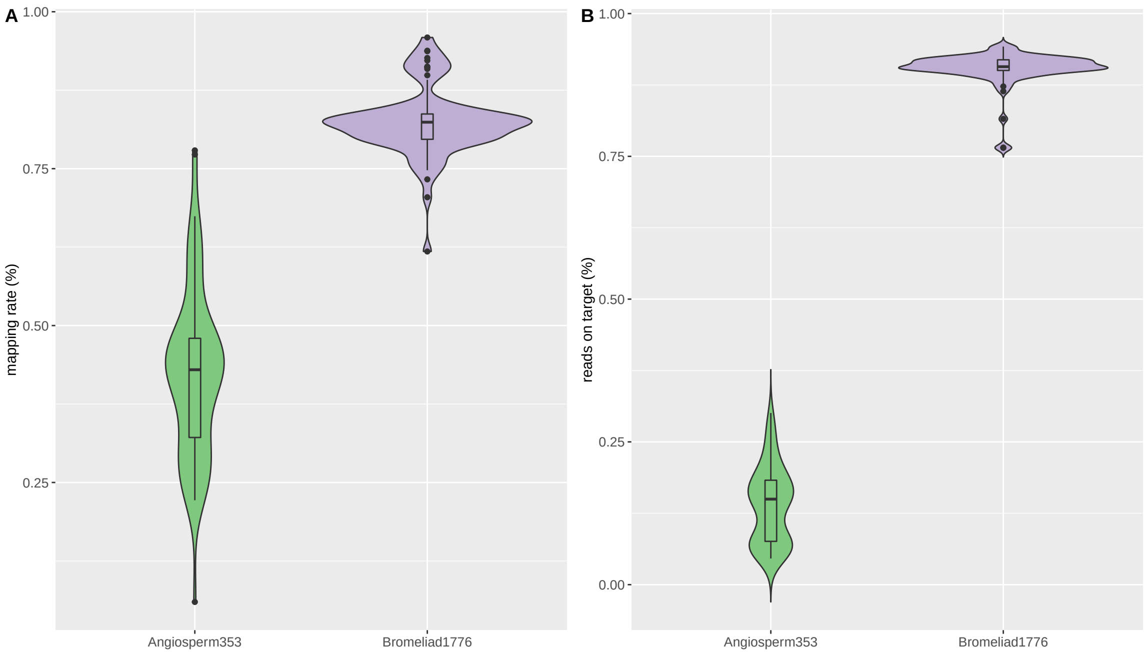


**Figure S2** Mapping rates (A) and percentage of reads matching bait sequences (B) for Bromeliad samples enriched with one of two bait sets: Angiosperms353 and Bromeliad1776. Reads were mapped against *A. comosus* reference for both bait sets. Targets were defined as bait locations and flanking 500 base-pairs. Bromeliad1776 targets were defined as the regions used for bait design and Angiosperms353 targets were defined as *A. comosus* orthologous regions matching the genes used for bait design.

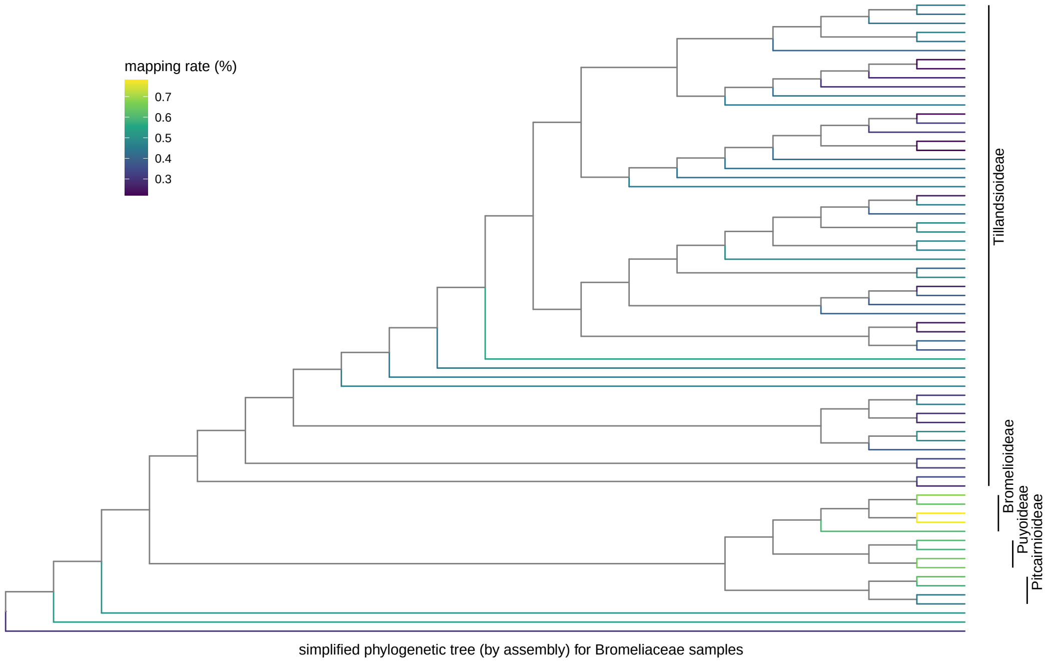


**Figure S3** A simplified phylogenetic tree, with branches colored according to read mapping percentage for samples enriched with Angiosperms353.

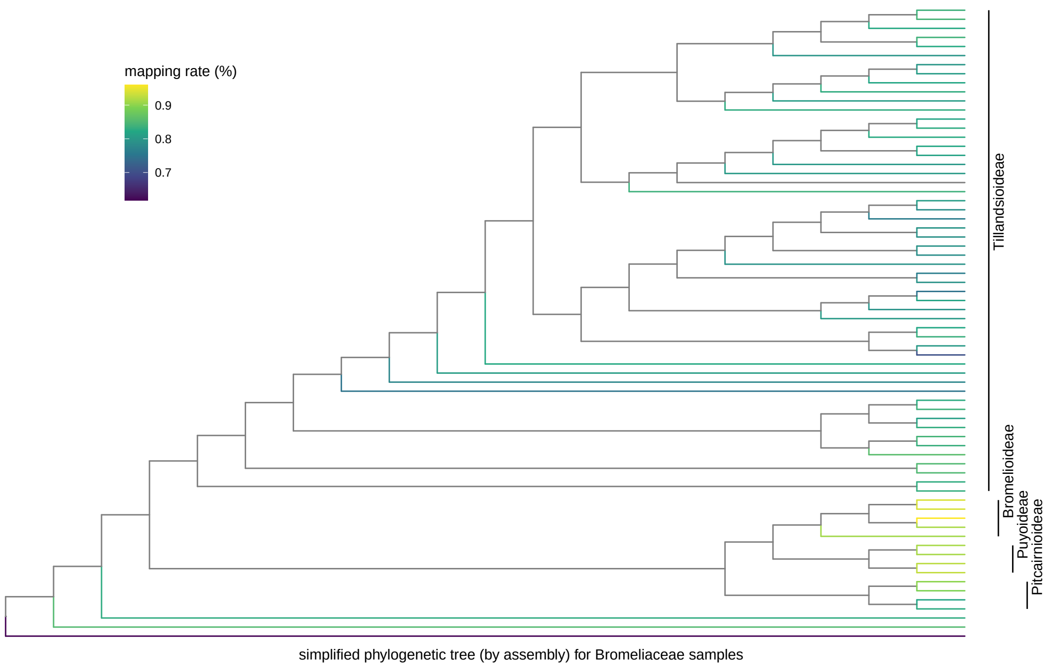


**Figure S4** A simplified phylogenetic tree, with branches colored according to read mapping percentage for samples enriched with Bromeliad1776.

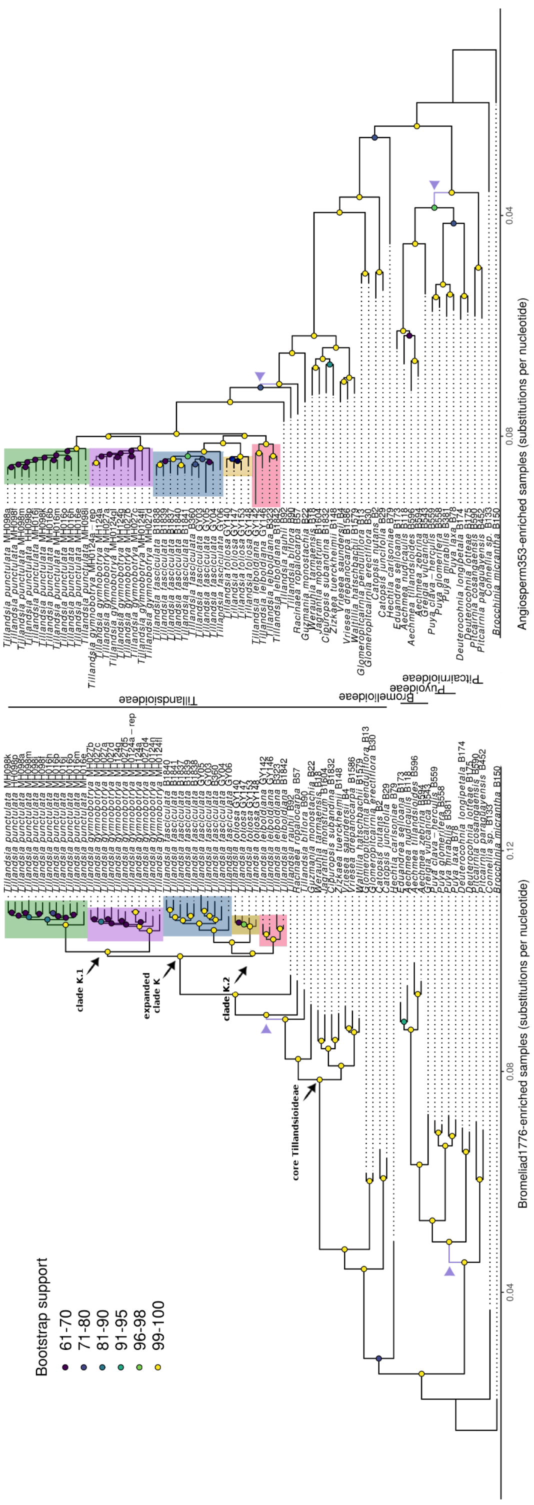
Bootstrap support
61-70
71-80
81-90
91-95
96-98
99-100
clade K.1
expanded clade K
clade K.2
core Tillandsioideae
0.04
0.08
0.12
Bromeliad1776-enriched samples (substitutions per nucleotide)
Tillandsioideae
Bromelioideae
Puyoideae
Pitcairnioideae
Angiosperm353-enriched samples (substitutions per nucleotide)

**Figure S5** Maximum-likelihood (ML) phylogenetic tree inferred with RAxML-NG, based on variants called for data sets enriched with Bromeliad1776 bait set (left) and Angiosperms353 bait set (right, flipped for mirroring). Branch lengths were calculated by number of substitutions per site. Internal nodes are marked and colored according to bootstrap support. Nodes which differed among trees are colored purple and have been marked by an arrow.

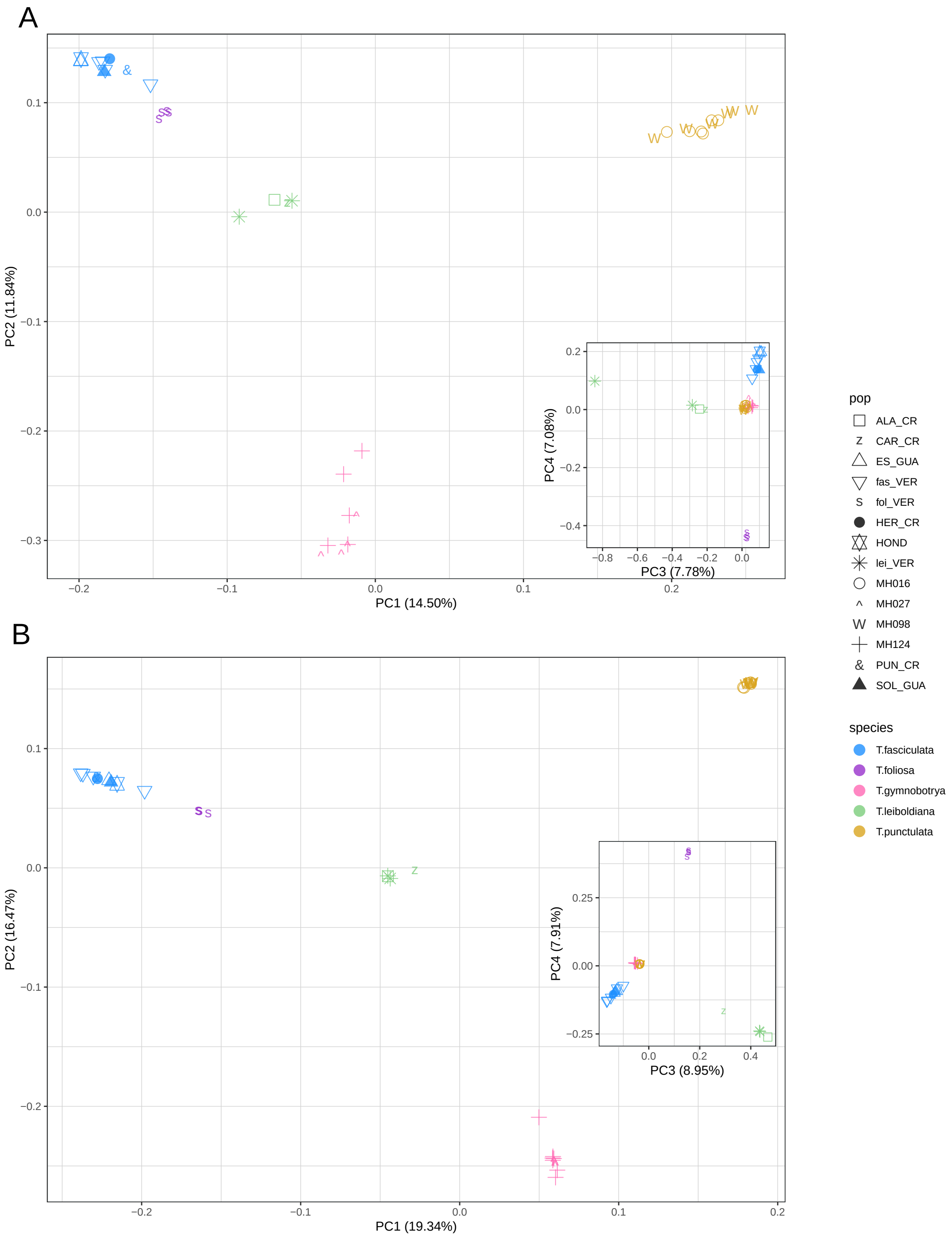
A
PC1 (14.50%)
PC2 (11.84%)
PC3 (7.78%)
PC4 (7.08%)
B
PC1 (19.34%)
PC2 (16.47%)
PC3 (8.95%)
PC4 (7.91%)
pop
ALA_CR
CAR_CR
ES_GUA
fas_VER
fol_VER
HER_CR
HOND
lei_VER
MH016
MH027
MH098
MH124
PUN_CR
SOL_GUA
species
T.fasciculata
T.foliosa
T.gymnobotrya
T.leiboldiana
T.punctulata

**Figure S6** Principal Component Analysis (PCA) plot for samples of *Tillandsia* subgenus *Tillandsia* enriched with two bait sets: A. Angiosperms353 (1,025 variants after LD-pruning) B. Bromeliad1776 (32,941 variants after LD-pruning). Colors indicate different species (following the scheme in Supporting Figure S6) and shapes represent different geographic origins (populations).

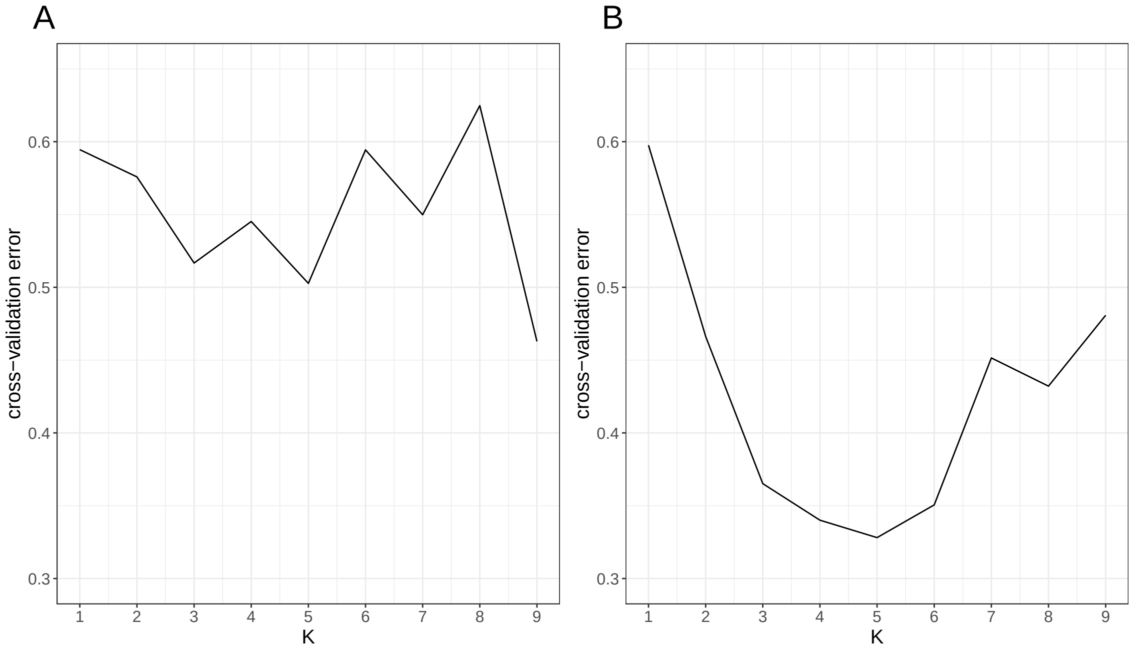


**Figure S7** Admixture cross-validation errors (top) detected for values of K between 2 and 9 for A. Angiosperms353 data set and B. Bromeliad1776 data set.
